# The BRD4-nucleosome interaction is enhanced modestly and non-selectively by histone acetylation

**DOI:** 10.1101/2025.05.29.656505

**Authors:** Lucien S. Lambrechts, Xavier J. Reid, Lucas Kambanis, Clement Luong, Erekle Kobakhidze, Andrea Daners, Yichen Zhong, Karishma Patel, Charlotte K. Franck, Hakimeh Moghaddas Sani, Cameron Taylor, Jason K. K. Low, Richard J. Payne, Joel P. Mackay

## Abstract

BRD4 regulates gene transcription in complex eukaryotes, in part through the binding of its tandem bromodomains to acetylated lysine residues found in histones and transcription factors. Despite pharmacological inhibition of these domains showing promise in preclinical studies, clinical trial data have been less encouraging so far. A stronger understanding of BRD4 biochemistry could provide a route to better outcomes. To advance on prior work, which has focused almost entirely on the binding of isolated bromodomains and acetylated peptides, we have sought the preferred nucleosomal binding partner of full-length BRD4. We demonstrate that BRD4 binds with sub-micromolar affinity to both unmodified nucleosomes and to DNA alone. In strong contrast to BRD4-peptide interactions, we also find that the affinity of BRD4 for nucleosomes is increased only 2–4-fold by histone acetylation and that this affinity has little dependence on the acetylation pattern. Despite this modest effect of acetylation, binding of BRD4 to acetyllysine in the nucleosome was more resistant to perturbation by mutation or small-molecule inhibition than BRD4-peptide interactions. Our work helps bridge the gap between cellular and prior in vitro work and provides clues to explain the in vivo chromatin occupancy profile of BRD4 and how it changes upon therapeutic inhibition.

**GRAPHICAL ABSTRACT:** 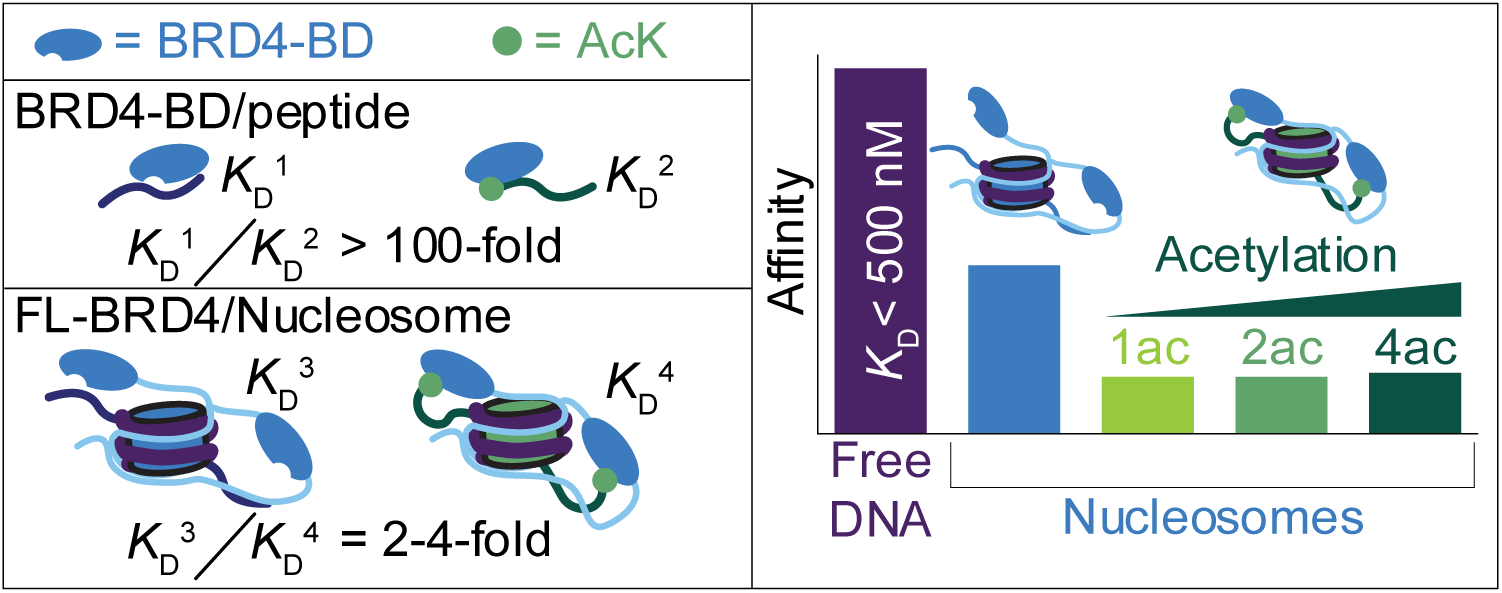

## INTRODUCTION

Bromodomain Extra-Terminal domain (BET) proteins are a set of widely and abundantly expressed transcriptional regulators that have homologues across eukaryotes (2–4). In mammals, the family comprises BRD2, BRD3, long and short isoforms of BRD4 (BRD4S and BRD4L), and the testis-specific BRDT. Both BRD2 and BRD4 are essential for normal development and all family members appear in general to act as transcriptional coactivators (5–7).

BET proteins feature two tandem bromodomains (BDs), which are well-established as readers of lysine acetylation (2) (**Figure 1A**). The conventional model for BET protein function centres on their bromodomain-dependent binding to acetylated histones or transcription factors. Most work has focused on interactions with acetylated histones – particularly histones H3 and H4 bearing lysine acetylation on their *N*-terminal tails (8,9). High levels of H3 and H4 acetylation are strongly associated with the promoters and enhancers of active genes and numerous genome-wide chromatin immunoprecipitation (ChIP-seq) experiments show that BET proteins are also enriched at these locations (10,11).

**Figure 1:**
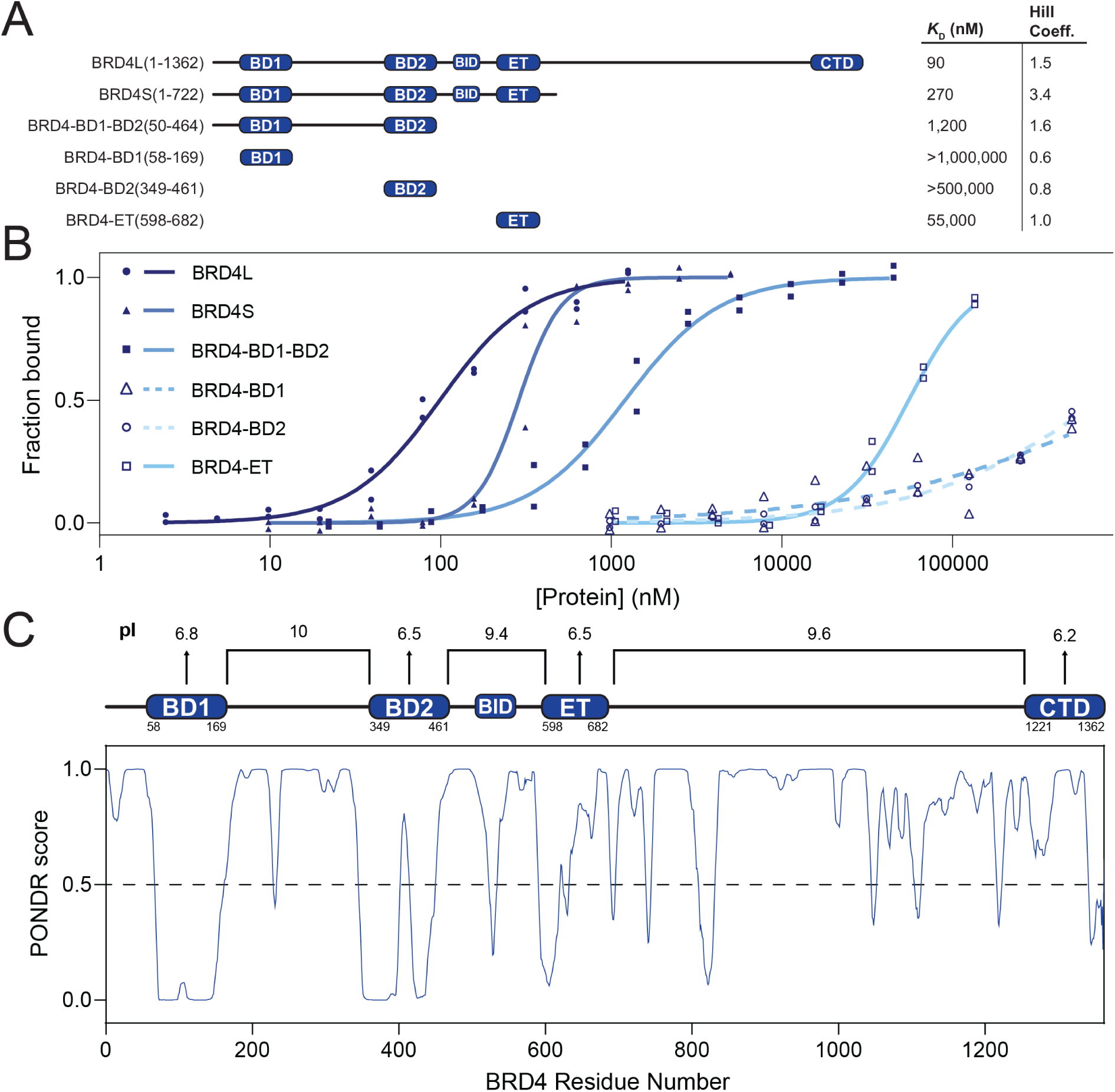
BRD4 binds naked DNA. **A.** Constructs of human BRD4 used in this study. Domains known/predicted to be ordered are labelled (BD1 = N-terminal bromodomain, BD2 = C-terminal bromodomain, BID = Basic-residue enriched interaction domain, ET = Extra-terminal domain, CTD = C-terminal domain) *K*D values derived from MST assays are also given, calculated from fits of titration data to the Hill equation. **B.** Technical duplicate MST assays for BRD4 isoforms/truncations binding to a Cy5-labelled 267-bp DNA (60N60). The SEM of all MST measurements is estimated to be 15% (see methods) **C.** Charge composition of ordered and disordered regions of BRD4 (*top*) and PONDR (1) disorder prediction (*bottom*).

A large body of *in vitro* work has corroborated these cell-based observations. Typically, these studies have interrogated binding of recombinant BET protein BDs to short histone peptides bearing a wide range of acetylation patterns (9,12–19). Measured *K*_D_ values for peptides bearing either single or multiple acetyllysines (AcKs) range from ∼10 to several hundred micromolar, and binding is strongly dependent on the acetylation pattern. For example, the N-terminal BD of BRD4 (BRD4-BD1) binds a histone H4 peptide bearing K5 and K8 acetylation (H4K5acK8ac) with a *K*D of 10 μM but the same BD binds an H4K8acK12ac peptide with a *K*D of 200 μM (18). Binding to the corresponding unacetylated peptide is undetectable, indicating selectivity of ∼100-fold for favoured acetylation patterns.

Concurrently, interactions between BET proteins and acetylated transcription factors (TFs) have also been demonstrated to have clear transcriptional outcomes. For example, acetylation of Nuclear Factor κB (NF-κΒ) drives gene expression during inflammation partially through an interaction with the BET proteins (20,21). Similarly, an interaction between acetylated GATA1 and BRD3 is required for GATA1 to bind and activate its target genes; this interaction simultaneously stabilizes the chromatin binding of BRD3 (22–24).

Structural analysis supports these findings: multiple three-dimensional (3D) structures of BET BDs bound to acetylated peptides derived from both histones and TFs have been reported (9,13,15,16,18,20,25,26). Altogether, these data suggest a preference for BD1 domains to bind diacetylated motifs with the consensus sequence AcK-XX-AcK (where X stands for any given amino acid), a motif that is found in histones H2A.Z and H4 and in multiple TFs. The preferences of BD2 domains are less clear but suggest weaker binding, predominantly to single acetylated motifs from histone H3 and H4.

Targeting of BET BDs with small molecules that compete for the AcK-binding pocket has shown significant therapeutic promise (17,27,28). However, despite promising pre-clinical data (29), however, BET inhibitors have not yet progressed past clinical trials (30). There remains therefore a need to deepen our understanding of the mechanisms by which BET proteins act.

Despite an abundance of studies describing the AcK-binding properties of BET-family proteins, almost all have focused on single or tandem BDs binding to short histone- or TF-derived peptides (9,12,13,16,17,19,20,22). In an effort to better recapitulate the behaviour of cellular BRD4, we have examined the binding of full-length BRD4 to recombinant nucleosomes with a range of architectures and acetylation patterns. We report that the binding of BRD4 to unmodified nucleosomes and DNA is significantly tighter than expected. Surprisingly, acetylation of nucleosomal histones increases the affinity for BRD4 by only 2–4 fold relative to unmodified nucleosomes and this effect is largely independent of the specific histone acetylation pattern. Furthermore, the ability of the BET inhibitor JQ1 to compete with an acetylated nucleosome for BRD4 binding is compromised. These data provide the first quantitative view of the nucleosome binding properties of BRD4 and suggest possible explanations for the observed cellular behaviour of BET proteins and their interactions with chromatin, transcription factors and small molecules

## MATERIAL AND METHODS

### Unmodified histone expression and purification

Unmodified histones were expressed and purified as described previously (31). Human histones H2A, H2A.Z, H2B and H3.1 encoded in pET28a plasmids were expressed in *Escherichia coli* BL21(DE3) cells overnight in ZYM-5052 auto-induction media at 37 °C with shaking at 180 rpm supplemented with 50 ng/μL kanamycin (32). H4 was expressed in the same host but in Lysogeny Broth (LB) at 37 °C and induced at an OD of 0.8 with 1 mM isopropyl ß-D-1-thiogalactopyranoside (IPTG) for 4 h. Cells expressing human histones were lysed by sonication in 50 mM Tris-HCl pH 7.5, 100 mM NaCl, 1 mM β-mercaptoethanol (BME) and clarified by centrifugation at 15,000 g for 30 min. The insoluble fraction was washed twice with the same lysis buffer containing 1% Triton X-100 and twice without. Inclusion bodies were extracted in 20 mM Tris-HCl pH 7.5, 6 M guanidinium HCl and 1 mM dithiothreitol (DTT)) with stirring overnight at 4 C. After clarification, filtered supernatants were injected onto a preparative Vydac protein and peptide C18 column (300-Å pore size, Catalogue No. 218TP1022) at a flow rate of 7 mL/min (20–70% acetonitrile over 40 min, 0.1% trifluoroacetic acid (TFA)). The fractions containing the target protein (as judged by liquid chromatography-mass spectrometry (LC-MS) analysis) were lyophilised and stored at –20 °C.

### Amber stop codon suppression for expression of acetylated H3 histones

mbAcKRS3-pCDFDuet1 plasmid was a gift from Andrea Cochran (Genentech) (33). Amber stop-codon mutations were induced in pET28a plasmids encoding histone H3.1 at nucleotides corresponding to H3K14 and to both H3K9 and K14 simultaneously. Amber mutant histones were expressed in *Escherichia coli* C321.ΔA cells co-transformed with mbAcKRS3-pCDFDuet1 in LB supplemented with 25 ng/μL kanamycin and 25 ng/μL spectinomycin at 37 °C with shaking at 180 rpm. Once an OD600 of 0.8 was reached, acetyllysine solution was added to a final concentration of 5 mM and expression was induced with 1 mM IPTG. Cells were grown for an additional 18 h. Cells expressing modified histone protein were lysed, and the inclusion bodies were extracted identically to unmodified histones. Extracted inclusion bodies were dialysed into 7 M urea, 20 mM NaOAc (pH 5.2) 200 mM NaCl, 5 mM BME, 1 mM EDTA buffer and loaded onto a 5 mL HiTrap SP HP column (Cytiva) pre-equilibrated in the same buffer. Protein was eluted over a gradient of 200 mM NaCl to 1 M NaCl over 30 min. Fractions containing the target protein (as judged by SDS-PAGE) were pooled and dialysed into Milli-Q water, freeze-dried and stored at -20 °C in sealed containers. Identity and site specific acetyllysine incorporation was verified with intact tandem mass spectrometry acquired on a Fusion tribid mass spectrometer (Thermo Fisher Scientific).

### Truncated histone expression and purification

Gene fragments corresponding to residues 23-127 of H2A.Z-A23C, 29-135 of H3.1-A29C-C96S-C110S, 15-102 of H4-A15C and 38-102 of H4-A38C were amplified by PCR and subcloned into a pET28a vector with an N-terminal His-SUMO tag using Gibson Assembly (34). SUMO-truncated histone fusions were expressed using the same autoinduction method as for unmodified histones. Cells were lysed either by sonication or by homogenisation with an Emulsiflex C3 homogeniser in 0.1 mM EDTA, 50 mM Tris, 500 mM NaCl, 0.05% v/v β-mercaptoethanol, 2 mM PMSF, 1X cOmplete™ Mini EDTA-free protease inhibitor, pH 7.5 and clarified by centrifugation at 15,000 g for 30 min. The insoluble fraction was washed twice with 50 mM Tris-HCl pH 7.5, 100 mM NaCl, 1 mM β-mercaptoethanol (BME) containing 0.5% w/v sodium deoxycholate and twice without. Inclusion bodies were extracted in 20 mM Tris-HCl pH 7.5, 6 M guanidinium HCl and 5 mM tris(2-carboxyethyl) phosphine (TCEP) with stirring overnight at 4 C. Extracted inclusion bodies were clarified by centrifugation at 15,000 g for 30 min and purified by incubating with nickel-nitrilotriacetic acid (Ni-NTA) Agarose resin for 1 h and elution with 6 M guanidinium HCl, 500 mM imidazole, 5 mM TCEP. Protein concentration was estimated by absorbance at 280 nm. SUMO-fusion proteins were refolded stepwise into 6 M urea, 150 mM NaCl, 100 mM L-arginine, 20 mM Tris-HCl pH 7.5, 7.5 mM sodium ascorbate, 5 mM TCEP for 2 h followed by two additional dialysis steps into the same buffer with 4 M and 2 M urea respectively. In the final dialysis step, proteins were diluted to approximately 1 mg/mL. SUMO tags were cleaved following the addition of 1 mg of ULP1 (pFGET19_Ulp1 was a gift from Hideo Iwai; Addgene plasmid #64697; http://n2t.net/addgene:64697; RRID: Addgene_64697) and stirred at 4 °C for 18 h. Cleaved protein was precipitated by the addition of 15% v/v trichloroacetic acid and centrifugation at 15,000 g for 30 min. Precipitated proteins were re-extracted in 20 mM Tris-HCl pH 7.5, 6 M guanidinium HCl, 7.5 mM sodium ascorbate, 5 mM TCEP and were purified in RP-HPLC using a preparative C8-Xbridge column (5 μm, 20 x 150 mm) with a gradient of 20-70% MeCN with 0.1% TFA over 40 min at a flow rate of 14 mL/min. Identity of the purified protein was confirmed in UPLC-MS and pooled fractions were lyophilised.

### Histone peptide synthesis

Histone peptides corresponding to residues 1-22 of H2A.Z, 1-28 of H3, 1-14 of H4 and 1-37 of H4 with different acetylation states were synthesised using a standard fluorenylmethyloxycarbonyl (Fmoc)- strategy solid-phase chemistry using a SYRO 1 automated synthesiser (Biotage). 2-chlorotrityl chloride (2-CTC) resin (1.22 mol/g loading) was swollen in dichloromethane (DCM) for 30 min. Fmoc-protected amino acid (Fmoc-Xaa-OH) (3 molar eq. relative to maximum loading) and *N,N*-diisopropylethylamine (DIPEA) (15 eq) in DCM (to make up 0.3 M of Fmoc-Xaa-OH) was added to the resin and agitated for 18 h at 25 ℃. Loading solution was ejected and the resin was washed with *N,N*-dimethylformamide (DMF) (5 × 3 mL), DCM (5 × 3 mL) and DMF (5 × 3 mL). Resin was treated with 4 mL of capping solution (17:2:1 v/v/v DCM: methanol (MeOH): DIPEA) for 1 h and were then washed with DMF (5 × 3 mL), DCM (5 × 3 mL) and DMF (5 × 3 mL). Solutions of Fmoc-Xaa-OH (0.3 M in DMF), Oxyma (0.33 M in DMF) and DIC (0.3 M in DMF) were prepared. Resin-Xaa was extended using a SYRO Automated Synthesiser. Fmoc deprotection was performed by shaking resin in 2 mL of 40% v/v piperidine in DMF twice for 10 min. Deprotection solution was extracted and resin was washed 4 times with 4 mL of DMF for 30 s. For coupling, peptide was shaken in a solution of Fmox-Xaa-OH (6 eq, 0.1 M), Oxyma (6.6 eq, 0.11 M), and DIC (6 eq., 0.1 M) in DMF for 45 min at 25 ℃. Coupling solution was extracted and resin was washed 4 times with 6 mL of DMF for 30 s. Capping was performed by shaking peptide in 2 mL capping solution (0.3 M Ac_2_O, 0.3 M DIPEA in DMF) for 3 min. Capping solution was extracted and resin was washed with 4 times with 6 mL of DMF for 30 s. After the final coupling step, peptides were Fmoc deprotected and then re-protected with a Di-tert-butyl decarbonate (Boc2O) protecting group by incubating resins for 1 h in 0.2 M Boc2O, 0.05 M DIPEA in DCM.

### Thioesterification of histone peptides

Extended peptide chains were treated with 5 mL of 30% v/v hexafluoro-isopropanol (HFIP) in DCM and agitated at room temperature for 1 h. Resin treatment was repeated. Cleaved peptide solution was dried with N2 and dissolved in 5 mL of DMF followed by the addition of 5 eq. DIPEA and 30 eq. ethyl-3-mercaptopropionate according to estimated resin loading. The reaction mix was cooled to -30 ℃ and a solution of benzotriazol-1-yloxytripyrrolidinophosphonium hexafluorophosphate (PyBOP) in DMF was added dropwise to 5 eq. The reaction was left for 3 h at -30 ℃ under nitrogen atmosphere. Solution was dried with N2 and treated with 5 mL cleavage cocktail of TFA: triisopropylsilane: water (90:5:5) for 4 h. Peptides were extracted with 9 volumes of diethyl ether and purified in preparative HPLC using a preparative C8-Xbridge column (5 μm, 20 × 150 mm) with a gradient of 0-40% MeCN (solvent B) and MQW (solvent A) with 0.1% TFA over 65 min at a flow rate of 14 mL/min. Identity of the purified peptide was confirmed in UPLC-MS and pooled fractions were lyophilised.

### Native chemical ligation for semisynthesis of acetylated H2A.Z, H3 and H4 histones

C-terminal histone fragments bearing an N-terminal cysteine were weighed out and dissolved to 2.5 mM in 1 M HEPES pH 7.2, 6 M guanidinium HCl, 200 mM mercaptophenylacetic acid, 100 mM TCEP. Two equivalents of the corresponding N-terminal thioester histone peptides were weighed out and dissolved in the mix. Reactions were incubated with shaking at 37 C for 18 h. Success of the ligation was confirmed in UPLC-MS and reactions were quenched with the addition of 1 volume of 1 M HEPES pH 7.2, 6 M guanidinium HCl, 200 mM TCEP. Product was purified in RP-HPLC using a semi-preparative Waters C4 symmetry column using a gradient of 20-70% (over 60 min) MeCN containing 0.1% TFA (buffer B) and a flow rate of 4 mL/min. Identity of the purified protein was confirmed in UPLC-MS and pooled fractions were lyophilised. Dried ligated histones were dissolved to 10 mg/mL in buffer degassed under argon atmosphere containing 1 M HEPES pH 7.2, 6 M guanidinium HCl, 200 mM TCEP and 40 mM reduced glutathione. A solution of 320 mM 2,2’-azobis[2-(2-imidazolin-2-yl)propane] dihydrochloride (VA-044) was made up in methanol and added to the dissolved histones to a final concentration of 32 mM VA-044 and 10% methanol. Desulfurisation mixes were irradiated at 254 nm for 30 min. Success of the reaction was confirmed by UPLC-MS and reactions were desalted by dialysis into Milli-Q water (MQW) over three two-hour steps and one overnight step. The resultant modified histones were lyophilised and stored at –20 °C.

### Histone octamer refolding

Histones were refolded as previously described (31). Histones were mixed in equal ratios of H3 and H4 and 1.5 molar equivalents of H2A/H2A.Z and H2B in unfolding buffer (20 mM Tris-HCl pH 7.5, 6 M guanidinium HCl and 1 mM DTT). A typical 45-nmol scale refolding was dissolved in 1 mL of unfolding buffer and dialysed into 20 mM Tris-HCl pH 7.5, 2 M NaCl, 1 mM EDTA over three dialysis steps including one overnight. Refolded octamers were clarified by centrifugation at 15,000 g for 5 min and loaded onto a Superdex 200 Increase 10/300 GL 24 mL column. Histone octamers eluted in a peak around 12-13 mL as verified by SDS-PAGE. Fractions from this peak were pooled and concentrated to 50 μL.

### Nucleosomal DNA preparation

DNA oligonucleotides were amplified by PCR from a pGEM plasmid containing a 601-positioning sequence. The PCR primers contained 5′ Cy5 modifications (Integrated DNA Technologies, Singapore) to install a fluorophore in the DNA sequence. The PCR products were first concentrated by ethanol precipitation and were redissolved in 1× TE buffer (10 mM Tris-HCl pH 8, 1 mM EDTA), and then purified in a phenol:chloroform:isoamyl alcohol (25:24:1) extraction, followed by a chloroform wash to remove residual phenol. After isopropanol precipitation and washing with 70% (v/v) ethanol, the DNA pellet was dissolved in 1× TE and separated on a 0.5× TBE 5% polyacrylamide gel. The DNA band with the desired size was cut from the gel and electroeluted into 0.5× TBS at room temperature. The final DNA product was concentrated by ethanol precipitation overnight and the resulting pellet was dissolved in 1× TE.

### Nucleosome reconstitution

Mononucleosomes were assembled by salt gradient dialysis using a double dialysis method (35). After mixing labelled octamer and DNA at a 1:0.95 molar ratio in 10 mM Tris-HCl pH 7.5, 2 M NaCl, 1 mM EDTA, the mixture was loaded into a small dialysis button, which was then placed into a dialysis bag containing 30 mL of 10 mM Tri-HCl (pH 7.5), 2 M NaCl, 1 mM EDTA and 0.1 mM DTT. The bag was then dialysed against 2 L of 1× TE containing 0.1 mM DTT overnight at room temperature. The next day, the dialysis button was dialysed further against 50 mM HEPES pH 7.5, 1 mM EDTA and 0.1 mM DTT. Content in the dialysis button was harvested, and the nucleosome quality was evaluated by 0.5× TBE 5% polyacrylamide gel electrophoresis at 150 V at 4 C.

### Expression and purification of GFP-BRD4L in HEK293 cells

HEK Expi293F^TM^ cells were grown to a density of 2 x 10 ^6^ cells/mL in Expi293^TM^ Expression medium (Thermo Fisher Scientific) at 37 °C and 5% CO_2_ with shaking at 130 rpm. A transfection mix containing 2 μg/mL of culture of pcDNA-GFP-BRD4L plasmid, 8 μg/mL of culture in 100 μL/mL of culture of PBS was added to the culture. Protein was expressed for 72 h and harvested with centrifugation at 500 g for 5 min. Cells were lysed by sonication in 20 mM Tris (pH 7.5), 500 mM NaCl, 0.1% v/v Triton X-100, 3 mM ATP, 3 mM MgCl2, 0.2 mM DTT, 2 × cOmplete EDTA-free protease inhibitor, 2 mM PMSF and clarified by centrifugation at 15,000 g for 30 min. Per 200 mL of cell culture, 400 μL of high capacity streptavidin beads were pre-equilibrated in 20 mM Tris (pH 7.5), 150 mM NaCl, 0.1% v/v Triton X-100 and incubated with 50 μg of His-SUMO-StrepTagII-anti-GFP nanobody for 4 h. Clarified lysate was incubated with the GFP-nanobody-coupled streptavidin beads for 16 h and eluted in 20 mM HEPES (pH 7.5) 150 mM NaCl, 50 mM Biotin. Protein purity and concentration was evaluated by SDS-PAGE and SYPRO-Ruby staining.

### Expression and purification of BRD4S and BRD4S mutants in E. coli

pQE80L plasmids encoding GST-3C-AVI-BRD4S fusions with different mutations were transformed into Rosetta 2(DE3) *Escherichia coli* cells and grown at 37 °C with shaking at 180 rpm until an OD600 of 0.8 was reached. Cultures were cooled to 18 °C and expression was induced with 0.25 mM IPTG. Protein was expressed for 16 h and harvested by centrifugation at 4,500 g for 25 min. Cells were lysed by sonication in 20 mM Tris (pH 7.5), 500 mM NaCl, 0.1% v/v Triton X-100, 0.2 mM DTT, 2 × cOmplete EDTA-free protease inhibitor, 2 mM PMSF and clarified by centrifugation at 15,000 g for 30 min. Clarified lysate was incubated with GSH-Sepharose for 4 h. Beads were washed with twice with lysis buffer and twice with 20 mM HEPES, 150 mM NaCl, 0.5 mM TCEP before overnight cleavage with HRV-3C protease. Cleaved protein was loaded onto a 5 mL HiTrap SP HP column (Cytiva) pre-equilibrated in the same buffer. Protein was eluted over a gradient of 150 mM NaCl to 1 M NaCl over 30 min. Fractions containing the target protein (as judged by SDS-PAGE) were pooled and salt concentration was calculated from the elution volume. Protein was concentrated and subsequently diluted to give a final salt concentration of 150 mM NaCl.

### Expression and purification of BRD4 bromodomains, ET domain and Tandem bromodomains in E. coli

GST-3C-AVI-BRD4-BD1, GST-3C-AVI-BRD4-BD2, GST-3C-AVI-BRD4-BD1-BD2 fusion genes encoded in pQE80L were expressed in *E. coli* BL21(DE3) cells and purified as previously described (36) with both GSH and Size-exclusion chromatography with a HiLoad 16/600 Superdex 75 SEC column. HA-GST-3C-BRD4-ET fusion gene encoded in pGEX was expressed in *E coli* BL21(DE3) cells identically to other BRD4 truncations and purified with only GSH affinity chromatography followed by on-beads cleavage of the GST with HRV-3C and elution of the desired protein.

### Microscale thermophoresis

A serial 1:1 dilution of BRD4 protein was prepared in 20 mM HEPES (pH 7.5), 150 mM NaCl with 2.5 μL of each concentration point. To each concentration point, 2.5 μL of a master mix was added to give a final volume of 5 μL containing 20 mM HEPES (pH 7.5), 75 mM NaCl 0.1 mg/mL bovine serum albumin, 0.8% glycerol and 20 nM of Cy5-labelled DNA or nucleosome. The mixtures were incubated on ice for 5 min and loaded into MST Monolith premium capillaries. Capillaries were inserted into a Monolith NT.115 MST instrument and scanned at 45% LED power for quality control. MST runs were carried out at 20% MST power for 15 s and 95% LED power. Fluorescence change was calculated from the average fluorescence count from -1–0 s before heating and 1.5–2.5s after heating. Scans of each capillary were run in triplicate with values for each titration calculated as an average for the triplicate scans. Each titration was repeated in at least two technical replicates defined as a new dilution series with the same protein and nucleosome prepared from the same DNA and octamer. Data was fitted to a Hill-Langmuir binding curve with a fixed analyte concentration using NanoTemper™ MO. Affinity Analysis software. Measurements were repeated on different unmodified 0N0 and H4K5acK8ac 0N0 nucleosome preparations at least 5 times. The SEM of these measurements was ∼15%; this value was used to estimate uncertainties for all MST titrations.

### Electrophoretic mobility shift assays

Unmodified Cy5 labelled 46N60 nucleosome was incubated with BRD4 isoforms and truncations in a mixture containing 50 nM nucleosome, 50 mM HEPES (pH 7.5), 50 mM NaCl and incubated on ice for 20 min. Sucrose was added to a final concentration of 5% (w/v) and titration points were loaded in 0.5 × TBE 5% polyacrylamide gels and run at 150 V for 1 h. Gels were imaged on a Typhoon FLA-9000 laser scanner.

### Surface-plasmon resonance

SPR measurements were taken on a Biacore T200 (Cytiva) instrument. WT-BRD4S or BRD4S-N140A-N433A was diluted to 0.5 μM in 10 mM sodium acetate (pH 5.5) buffer and immobilised on a Series S Sensor chip CM5 with *N*-hydoxysuccinamide/*N*-(3-dimethylaminopropyl)-*N’*-ethylcarbodiimide hydrochloride (NHS/EDC)-based amine coupling at 25 °C to a target density of 2500-5000 RU. Flow cells were blocked with an injection of 1 M ethanolamine following immobilisation. Experiments were conducted at 4 °C in 150 mM NaCl, 20 mM HEPES (pH 7.5), 0.5 mM TCEP, 0.1% v/v Tween-20 with a flow rate of 50 μL/min. H4 peptides and JQ1 were injected using a multi-cycle kinetics mode with a contact time of 60s and a dissociation time of 300s. Data was analysed using Biacore T200 Evaluation software and fit to a 1:1 Langmuir binding isotherm. Where saturation was not reached, the Rmax was set using Rmax values derived from titration of an H4K5acK8ac peptide over WT-BRD4S.

### Fluorescence polarisation

Fluorescence polarisation assays were performed in a Costar 384-well flat bottom black polystyrene assay plate. Titration points were made up in 20 μLwith BRD4S (top concentration 1 μΜ), 20 nM FITC-JQ1 (purchased from bio-techne, catalogue number 7722/2), 75 mM NaCl, 20 mM HEPES (pH 7.5) and where indicated, 50 nM of either unmodified or H4K5acK8ac 0N0 nucleosome. Reactions were incubated at RT for 10 min and fluorescence polarisation was measured using a PHERAstar plate reader following excitation at 485 nm. Data were fitted to a 1:1 Langmuir binding isotherm using GraphPad Prism.

### Extraction and digestion of endogenous mononucleosomes for HEK293 cells

Suspension-adapted HEK Expi293F™ cells (100 mL) were grown in Expi293™ Expression Medium at 37 °C, 5% CO_2_ to a density of roughly 5 × 10^6^ cells/mL. The cells were spun down at 300 × *g* and 4 °C for 5 min, and the cell pellets were washed twice with phosphate buffered saline (PBS). Cell pellets were lysed in PBS supplemented with 0.3% (v/v) Triton-X-100, 2× cOmplete EDTA-free protease inhibitor, and 1× home-made phosphatase inhibitor cocktail (2 mM NaF, 2 mM Na₃VO₄, 2 mM β-glycerophosphate, 2 mM Na4P2O7), by inverting gently on ice for 10 min. Nuclei were pelleted by centrifugation at 680 × *g* and 4 °C for 5 min. Nuclei were washed twice with PBS supplemented with 2× cOmplete EDTA-free protease inhibitor and 1× home-made phosphatase inhibitor cocktail. Nuclei were resuspended in digestion buffer (10 mM HEPES pH 7.6, 20 mM NaCl, 1.5 mM MgCl2, 0.5 mM EGTA, 10% glycerol (v/v), 2× cOmplete EDTA-free protease inhibitor, 1× home-made phosphatase inhibitor cocktail, 1 mM DTT), and CaCl2 was added to a final concentration of 2 mM.

Chromatin was then digested into mononucleosomes with the addition of micrococcal nuclease (MNase) (New England BioLabs, Beverly, MA). MNase was added to nuclei at 4000 GU or 0.4 U per ∼120 million cells and incubated at 26 °C for 25 min. To stop the MNase digestion, EGTA was added at a final concentration of 10 mM. Digested nuclei were spun down at 20,000 × *g* and 4 °C for 30 min to remove any insoluble, undigested heterochromatin. The extent of MNase digestion was checked by phenol-chloroform DNA extraction followed by agarose gel electrophoresis.

The nucleosomes were further purified from other nuclear proteins by sucrose gradient ultracentrifugation. The MNase digested supernatant was loaded onto an 11-mL sucrose gradient made from a light buffer containing 10 mM HEPES pH 7.6, 20 mM NaCl, 1.5 mM MgCl2, 0.5 mM EDTA, 2× cOmplete EDTA-free protease inhibitor, 1× home-made phosphatase inhibitor cocktail and 7% sucrose (w/v), and a heavy buffer containing the same buffer components but with the addition of 30% sucrose (w/v). The sample was then spun at 34,000 rpm and 4 °C for 16 h using a SW41TI rotor. Fractions (200 µL) were collected manually from the top of the sucrose gradient, and fractions containing nucleosomes were confirmed by a combination of SDS-PAGE, native-PAGE and UV absorbance at 260 nm. Fractions containing nucleosomes were pooled, dialysed into 10 mM HEPES pH 7.6, 1.5 mM MgCl2, 0.5 mM EDTA using 10-kDa MWCO dialysis tubing, and then snap frozen and stored at –80 °C.

### Pulldown of endogenous nucleosomes with GFP-BRD4L

HEK Expi293F™ cells were transfected with GFP-BRD4L and lysed as previously described. Clarified lysates were incubated with streptavidin beads coupled to GFP-nanobody for 4 hand washed three times with 50 mM HEPES (pH 7.5), 500 mM NaCl, 0.1% v/v Triton X-100 3 mM ATP, 3 mM MgCl2, 0.2 mM DTT, twice with 50 mM HEPES (pH 7.5), 150 mM NaCl, 0.5% v/v IGEPAL CA630, 0.2 mM DTT and twice with mononucleosome wash buffer (10 mM HEPES (pH 7.6), 50 mM NaCl, 1.5 mM MgCl2, 0.5 mM EDTA). Endogenous mononucleosome at a final concentration of 1 μM was added to the beads and the final buffer concentration was made up to 50 mM NaCl, 0.2 mM DTT and incubated overnight at 4 °C. Supernatant was removed and beads were washed 5 times with mononucleosome wash buffer. Mononucleosomes were eluted in 50 mM HEPES (pH 7.5), 150 mM NaCl, 50 mM biotin and dialysed into mononucleosome wash buffer before lyophilisation.

### Mass spectrometry

HEK293-NCPs were prepared for mass spectrometry analysis as described previously (42,43). Nucleosomes were first propionylated by adding propionylation reagent (1:3 propionic anhydride:acetonitrile v/v) at a ratio of 1:4 (propionylation reagent:sample volume). The pH was checked and adjusted to 8.0 using 28% (w/v) ammonium hydroxide solution if necessary. The samples were incubated at room temperature for 15 min. This process was repeated a second time and then the samples were freeze-dried. After freeze drying, a second double round of propionylation was carried out to ensure >95% sample propionylation.

The samples were resuspended in 50 mM NH_4_HCO_3_ to a final concentration of ∼ 1 µg/µL and the pH was adjusted to 8.0. Trypsin was added at a ratio of 1:10 (w/w) and the samples were incubated overnight at 37 °C. Trypsin digestion was stopped by snap-freezing samples at –80 °C. Samples were freeze-dried and then reconstituted in 50 mM NH_4_HCO_3_ for another double round of propionylation as described above.

Detergents were then removed from the sample using the SP3 bead protocol (37). Samples were resuspended in 50 µL 2% (v/v) formic acid (FA), 950 µL 100% acetonitrile and SP3 beads (20 µL, 50 mg/mL) were added to the resuspended sample. The sample was incubated on a rotary tube mixer for 8 min at room temperature. The beads were washed once with 100% acetonitrile and then air-dried for 1 min. Peptides were eluted twice from the beads with 2% (v/v) dimethyl sulfoxide (DMSO) and then centrifuged at 17000 × *g* to remove any remaining beads. The supernatant containing peptides was retained and freeze dried.

Samples were resuspended in 2% (v/v) FA. Samples were then desalted using C18 columns (Waters) equilibrated with 80% (v/v) acetonitrile, 0.1% (v/v) FA. The columns were then washed twice with 4% (v/v) acetonitrile, 0.1% (v/v) FA. Samples were then applied to the columns and washed twice with 4% (v/v) acetonitrile, 0.1% (v/v) FA before eluting twice with 50% (v/v) acetonitrile, 0.1% (v/v) FA. The elutions were then pooled and freeze dried.

For LC-MS/MS, ∼0.5–1 μg of propionylated histones in loading buffer (4% (v/v) acetonitrile, 0.1% (v/v) FA) were injected onto a 30 cm × 75 μm inner diameter column packed in-house with 1.9-μm C18AQ particles (Dr Maisch GmbH HPLC) using a Dionex Ultimate 3000 nanoflow UHPLC. Peptides were separated using a linear gradient of 5–35% buffer B over 130 min at 300 nL/min at 55 °C (buffer

A consisted of 0.1% (v/v) FA and buffer B consisted of 80% (v/v) acetonitrile and 0.1% (v/v) FA). All MS analyses were performed using a Q-Exactive HFX mass spectrometer. For DDA: after each full-scan MS1 (R = 120,000 at 200 *m/z*, 300–1600 *m/z*; 3 × 10^6^ AGC; 110 ms max injection time), up to 10 most abundant precursor ions were selected for MS/MS (*R* = 45,000 at 200 *m/z*; 2 × 10^5^ AGC; 86 ms max injection time; 30 normalised collision energy; peptide match preferred; exclude isotopes; 1.3 *m/z* isolation window; minimum charge state of +2; dynamic exclusion of 15 s). This resulted in a duty cycle of ∼1.3 s. For DIA: after each full-scan MS1 (*R* = 60,000 at 200 *m/z* (300–1600 *m/z*; 3 × 10^6^ AGC; 100 ms max injection time), 54 × 10 *m/z* isolations windows (loop count = 27) in the 390–930 *m/z* range were sequentially isolated and subjected to MS/MS (*R* = 15,000 at 200 *m/z*, 5 × 10^5^ AGC; 22 ms max injection time; 30 normalised collision energy). 10 *m/z* isolation window placements were optimised in Skyline (44) to result in an inclusion list starting at 395.4296 *m/z* with increments of 10.00455 *m/z*. This resulted in a duty cycle of ∼2.2 s.

### Mass spectrometry data analysis

Database searches were performed using Mascot v2.8. Spectra were searched against the human histone database using a precursor-ion and product-ion mass tolerance of ± 20 ppm and ± 0.02 Da, respectively. The enzyme was specified as ArgC with 1 missed cleavage allowed. Variable modifications were set as follows: acetyl(K), propionyl(K), monomethyl + propionyl(K), dimethyl(K), trimethyl(K), propionyl(N-term), oxidation(M), and carbamidomethyl(C).

All DIA data were processed using Skyline (38). Reference spectral libraries were built in Skyline with .dat files using the BiblioSpec algorithm (39). A False Discovery Rate (FDR) of 5% was set and a reverse decoy database was generated using Skyline.

Precursor and product ion extracted ion chromatograms were generated using extraction windows that were two-fold the full-width at half maximum for both MS1 and MS2 filtering. Ion-match tolerance was set to 0.055 *m/z*. For MS1 filtering, the first three isotopic peaks with charges +2 to +4 were included while for MS2, *b-* and *y-*type fragments ions with charges +1 to +3 were considered.

Peaks were identified and assigned based on (i) the dot product between peptide precursor ion isotope distribution intensities and theoretical intensities (idotp ≥ 0.9), (ii) the retention times of identified peptides based on Mascot searches and relative retention times based on hydrophobicity of PTMs (40), and (iii) unique MS2 fragment ions. Ultimately, three precursor ions (M, M + 1, and M + 2) were used for quantitation.

To quantify histone modification proportions, the resulting data were normalised by calculating peptide intensity as a proportion of the total intensity of a peptide family. A peptide family is defined as a group of peptides spanning the same residues within the histone H3 or H4 proteins but bearing different modifications. For instance, H3 residues 9–17 (KSTGGKAPR), which contains lysine K9 and K14, is a peptide family containing 10 peptides: H3 K9unK14un, K9me1K14un, K9me2K14un, K9me3K14un, K9acK14un, K9unK14ac, K9me1K14ac, K9me2K14ac, K9me3K14ac, and K9acK14ac. The peak area of each individual peptide was divided by the sum of peak areas of all peptides within the same peptide family to get a relative proportion for each peptide within that family. The cumulative proportions for each modification at a specific residue were then calculated by summing all peptides bearing the modification.

Fold changes in the proportion of modified histone peptides after pulldown with BRD4 were determined by comparing to the input endogenous nucleosome library described in Reid et al. (in review). Mass spectrometry data for the input library, which were used to calculate peptide proportions obtained in the pulldown are available from PRIDE (41) partner repository with the dataset identifier PXD064241.

## RESULTS

### BRD4 binds DNA with nanomolar affinity

We first assessed the ability of BRD4 to bind naked DNA. We prepared an oligonucleotide comprising the 147-base-pair (bp) Widom positioning sequence with 60 bp of flanking DNA on each side (60N60) and bearing a Cy5 fluorophore on the 5’ end of one strand (**Supplementary Figure 1**). This DNA was titrated in microscale thermophoresis (MST) experiments with a range of BRD4 truncations (**Figures 1A, B**). The full-length long isoform of BRD4 (BRD4L), overexpressed and purified from HEK293 cells (**Supplementary Figure 2**), bound 60N60 with a *K*_D_ of 90 nM. Fits were performed using the Hill equation, based on the improved fit quality compared to a simple 1:1 binding isotherm. The two equations yielded the same *K*D values, but the Hill equation additionally provides a measure of the cooperativity of binding, under the assumption that more than one binding event is taking place. The Hill coefficient for BRD4L was 1.5, suggesting weakly cooperative binding of multiple BRD4L molecules to the DNA.

The short isoform of BRD4 (BRD4S, residues 1–722), which was expressed in *E. coli*, bound DNA with a *K*D of 280 nM and a Hill coefficient of 3.4, suggesting that multiple copies of BRD4S bind cooperatively to DNA. Truncation of BRD4S immediately after the second bromodomain, the so-called tandem bromodomain construct (50–464, BRD4-BD1-BD2) further reduced the affinity by 4-fold to 1.2 μM. The individual domains of BRD4 all bound 60N60 DNA with substantially lower affinity (*K*D values of >1 mM, >500 μM and 55 μM for BD1, BD2 and ET, respectively).

The BD1, BD2, ET and CTD regions in BRD4 have near neutral predicted pI values, whereas the regions linking these domains are strongly basic in nature (**Figure 1C, Supplementary Figure 3**). The MST data indicate that these latter regions, which are predicted to contain significant amounts of disorder (**Figure 1C**), contribute significantly to DNA binding, perhaps through the formation of ‘fuzzy’ electrostatic interactions with the DNA backbone.

### BRD4 binds unmodified nucleosomes with and without flanking DNA

We next tested the binding of BRD4 to nucleosomes assembled from recombinant human histones and the core 147-bp Widom sequence with 30 base pairs of flanking DNA either side (30N30, **Supplementary Figures 1, 4**). Both BRD4L and BRD4S bound to this 30N30 nucleosome approximately 2-fold tighter than to 60N60 DNA alone (*K*D = 44 and 150 nM, respectively; **Figures 2A, B**).

**Figure 2:**
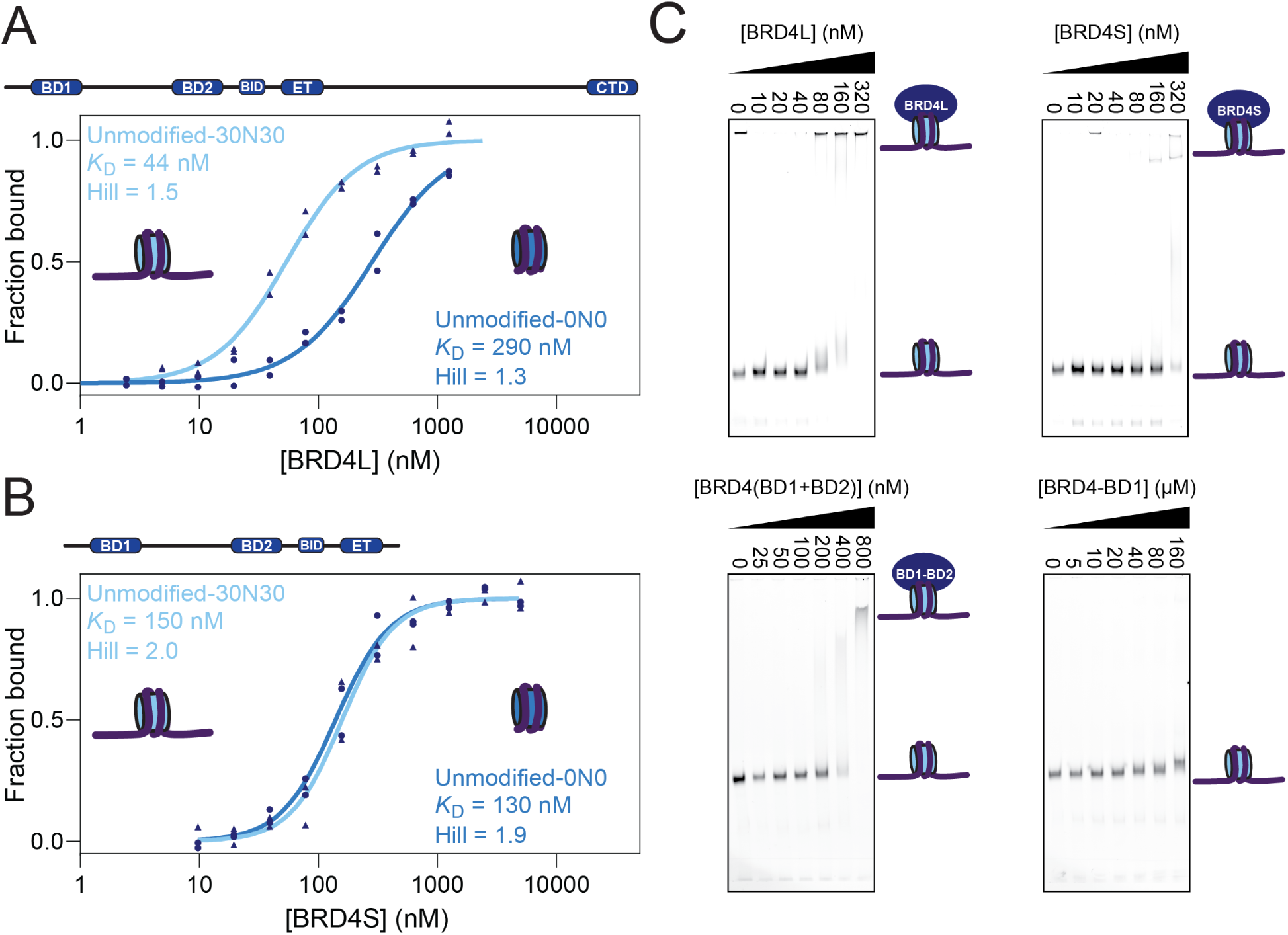
BRD4 binds unmodified nucleosomes. **A.** Technical duplicate MST assays for BRD4L binding to either 30N30 or 0N0 Cy5-labelled nucleosomes. **B.** Technical duplicate MST assays for BRD4S binding to either 30N30 or 0N0 Cy5-labelled nucleosomes. The SEM of all MST measurements is estimated to be 15% (see methods) **C.** EMSAs for BRD4 isoforms/truncations binding to 46N60 Cy5-labelled nucleosomes.

To corroborate these observations, we carried out Electrophoretic Mobility Shift Assays (EMSAs) with different BRD4 truncations and nucleosomes with flanking DNA (46N60) (**Figure 2C**). For BRD4L and BRD4S, the shifts in nucleosome migration in the gel occurred at concentrations similar to the *K*D’s measured by MST. An EMSA of BRD4-BD1-BD2 revealed binding with a *K*D of ∼400 nM, whereas BRD4-BD1 interacted only marginally with the nucleosome (*K*D >200 μM).

We then asked whether BRD4 could bind to just the nucleosome core particle. We constructed a nucleosome with just the 147-bp Widom sequence (0N0, no flanking DNA) and assessed its ability to bind BRD4L and BRD4S. We saw that while BRD4S bound the core particle with almost identical affinity to the 30N30 nucleosome, BRD4L bound around 3-fold weaker to nucleosomes without flanking DNA than to naked DNA with an affinity of 290 nM (*K*D = 290 nM, **Figures 2A, B**).

Together, these data indicate that BRD4 binds nucleosomes with only 2-fold tighter affinity than DNA alone, and that the BRD4L C-terminal region is involved in binding the flanking regions of the nucleosome.

### The nucleosome binding activity of BRD4 is weakly sensitive to histone lysine acetylation – and only in the absence of flanking DNA

As noted above, numerous studies have described the strong affinity and high selectivity of BET-BD1 domains for peptides containing diacetylated AcK-XX-AcK motifs (13), perhaps most prominently histone H4K5acK8ac (9,18). We therefore assembled a recombinant 30N30 nucleosome bearing the H4K5acK8ac mark (**Supplementary Figures 4, 5**) and measured the interaction of this species with BRD4. Unexpectedly, neither BRD4L nor BRD4S displayed any preference for acetylated over unmodified nucleosomes (**Figures 3A, B**).

**Figure 3:**
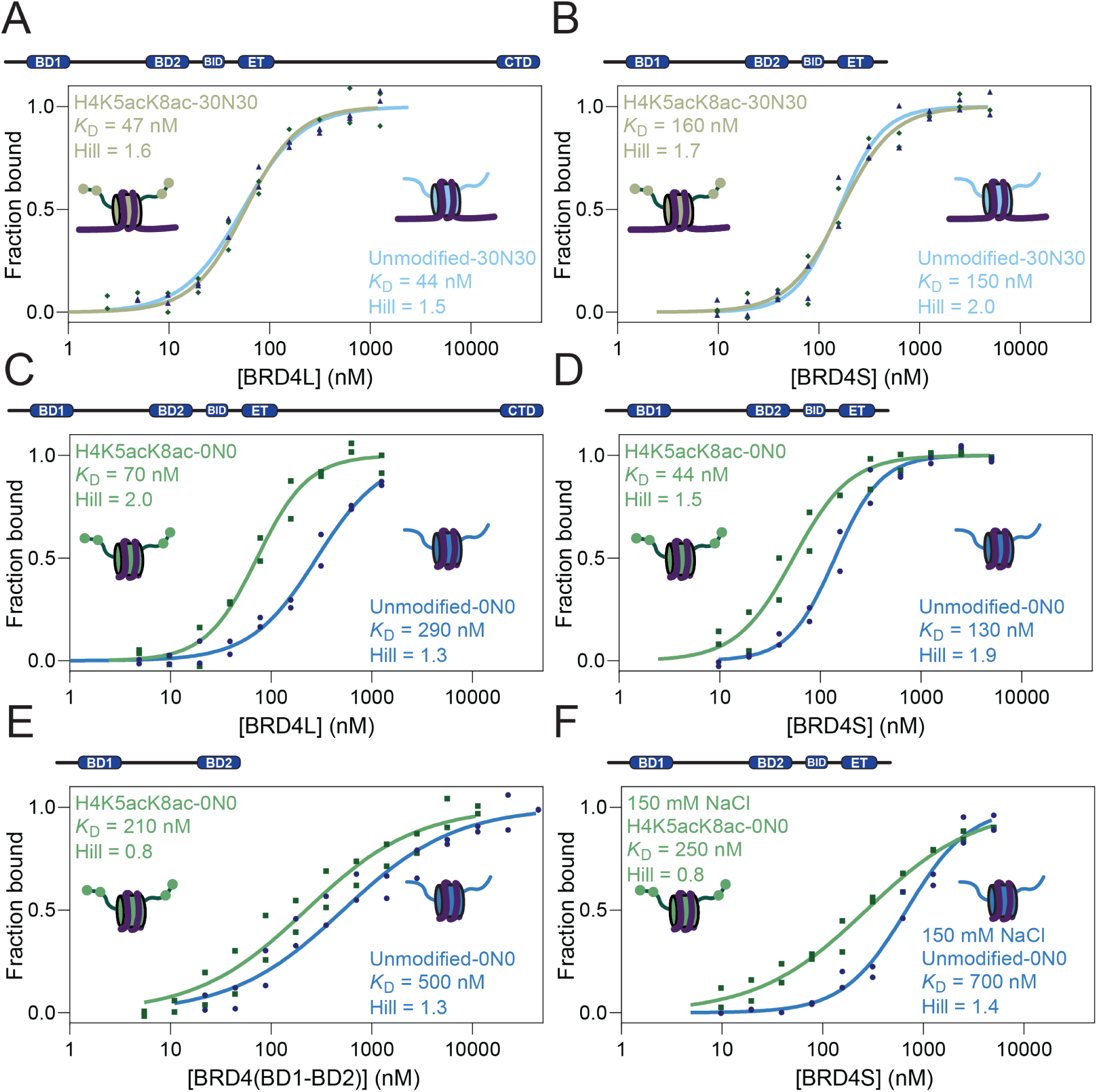
BRD4-nucleosome interactions are weakly dependent on histone acetylation. **A.** MST assays (technical duplicates) for BRD4L binding to either unmodified or H4K5acK8ac 30N30 Cy5-labelled nucleosomes. **B.** Duplicate MST assays (technical duplicates) for BRD4S binding to either unmodified or H4K5acK8ac 30N30 Cy5-labelled nucleosomes. **C.** MST assays (technical duplicates) for BRD4L binding to either unmodified or H4K5acK8ac 0N0 Cy5-labelled nucleosomes. **D.** MST assays (technical duplicates) for BRD4S binding to either unmodified or H4K5acK8ac 0N0 Cy5-labelled nucleosomes. **E.** MST assays (technical duplicates) for BRD4-(BD1-BD2) binding to either unmodified or H4K5acK8ac 0N0 Cy5-labelled nucleosomes. **F.** MST assays (technical duplicates) for BRD4S binding to either unmodified or H4K5acK8ac 0N0 Cy5-labelled nucleosomes in 150 mM NaCl. The SEM of all MST measurements is estimated to be 15% (see methods).

To assess whether the presence of flanking DNA somehow impacts access of the BDs to the H4 tail, we repeated the measurements with 0N0 nucleosomes. **Figure 3C–E** shows that, in this context, H4 di-acetylation enhanced binding of both BRD4S and BRD4-BD1-BD2 by 2–3-fold (*K*_D_ = 44 nM and 210 nM, respectively) and enhanced binding of BRD4L by 4-fold (*K*D = 70 nM). These acetylation-induced increases in affinity are robust but substantially smaller than previously described for isolated BET BDs binding histone peptides.

To this point, titrations had been carried out at 75 mM NaCl to preserve nucleosome stability. Given the significant interactions observed with naked DNA and the weak preference for acetylation, we repeated the titration of BRD4S into unmodified and H4K5acK8ac nucleosomes at 150 mM NaCl. Under these conditions, the affinity for both nucleosomes decreased by ∼5-fold, meaning that the preference for acetylation remained ∼3-fold (**Figure 3F**). Because the preference for acetylation was unchanged and that nucleosomes are more stable at the lower salt concentration, we continued to use 75 mM NaCl for our assays.

### BRD4 prefers acetylated nucleosomes when challenged with a native nucleosome library

Given the unexpectedly low sensitivity of BRD4 when binding H4 diacetylated nucleosomes, we conducted an unbiased screen with native nucleosomes to determine the nucleosome-binding preferences of BRD4. We first prepared a library of endogenous mononucleosomes by extracting chromatin from HEK293 cells and digesting with micrococcal nuclease (**Figure 4A**). We used mass spectrometry to determine the relative abundances of lysine acetylation and methylation on the N-terminal histone tails that could be detected (*i.e.*, those of H3, H3.3, and H4) in the library. Following lysine propionylation and trypsinisation, peptides corresponding to H3(3–8), H3(9–17), H3(18–26), H3.1/2(27–40), H3.3(27–40) and H4(4–17) were observable, and the relative abundances of distinct post-translational modification states were quantified by dividing intensities for each modification state by the total intensity for all peptides observed with a particular sequence.

**Figure 4:**
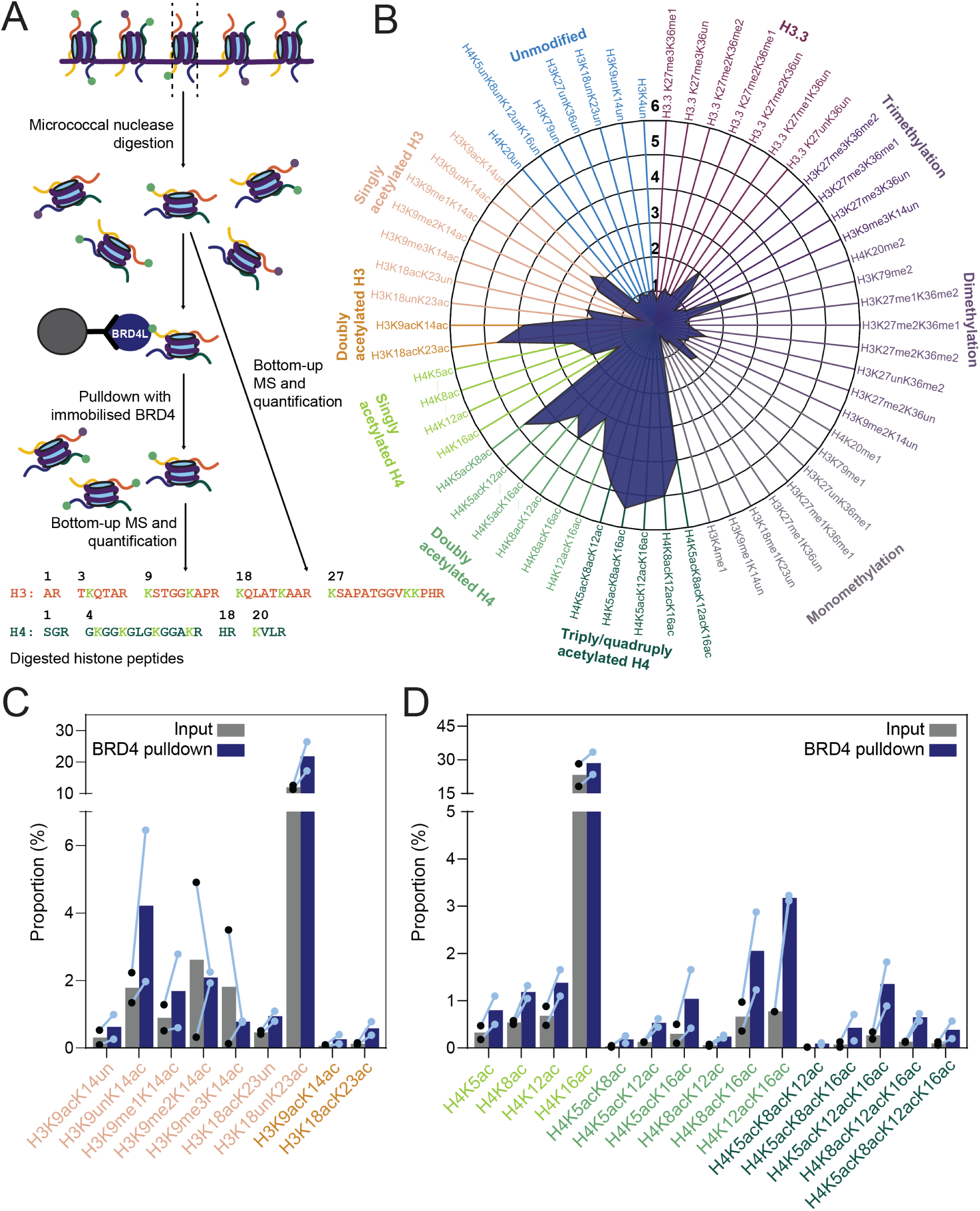
Modification preferences of BRD4-nucleosome pulldown. **A.** Schematic of method used to determine the histone PTM preferences of BRD4L. Mononucleosomes were extracted with micrococcal nuclease digestion and PTM abundance was quantified by bottom-up mass spectrometry. Streptavidin beads (pre-incubated with Strep-tagged anti-GFP nanobody and GFP-BRD4) were incubated with the nucleosome library, eluted and subjected to mass spectrometry. **B.** Radial plot outlining the mean fold changes in histone PTM distribution in the nucleosome library following capture by BRD4L in two biological replicates. **C.** Bar plot of proportion of H3-acetylated states in the nucleosome library before and after capture by BRD4. **D.** Bar plot of proportion of H4-acetylated states in the nucleosome library before and after capture by BRD4.

We next immobilised GFP-tagged BRD4L on streptavidin beads pre-incubated with Strep-tagged α-GFP nanobody and incubated the beads with our nucleosome library (**Figure 4A**). The proportions of each histone modification state were measured in the BRD4L-bound nucleosomes and compared to the input library. **Figure 4B** shows that, with the exception of H4K16ac, all acetylation states were enriched by at least two-fold. A five-fold increase was observed for nucleosomes tri-acetylated on H4 and a four-fold increase for nucleosomes di-acetylated on H3 or H4. Monoacetylated H3 and H4 nucleosomes were enriched by approximately two-fold. In contrast, methyllysine-containing sequences were not enriched, with the exception of H4K20me2. We note that the initial abundance of distinct acetylation marks was variable; for example, H4K16ac increased from 23% to 29% while H4K5acK8ac increased from 0.04% to 0.18%, which could contribute to variance in the measured fold change. (**Figures 4C, D, Supplementary Table 1**).

Overall, the data suggest that BRD4L prefers binding nucleosomes containing H3 and H4 acetylation, though we note that the method is blind to the modification state of other histones in the same nucleosome. For example, although we detect enrichment of H4 acetylation, we cannot determine whether or not the enriched H3-acetylated nucleosomes also contain H4 acetylation.

### BRD4 recognizes histone acetylation promiscuously

To explore the native nucleosome enrichment data in a defined system, we constructed a library of acetylated recombinant 0N0 nucleosomes and probed their binding to BRD4S by MST (**Figures 5A, B, Supplementary Figures 4-7**). We included H2A.Z in the library because its *N*-terminal tail can be acetylated and because two of the AcKs form a AcK-XX-AcK motif (K4+K7). Unexpectedly, all tested combinations of acetylation marks increased the affinity for BRD4S by a similar amount: approximately 2–4-fold. The greatest increases were observed for H3K14ac and H4K5ac. Acetylation of H2A.Z did little to enhance binding beyond the 2–3-fold increase already observed upon incorporation of unacetylated H2A.Z. Acetylation of multiple histone tails including H2A.Z and H4 together and H3 and H4 together also showed no further increases in affinity, despite each of these modifications individually enhancing affinity.

**Figure 5:**
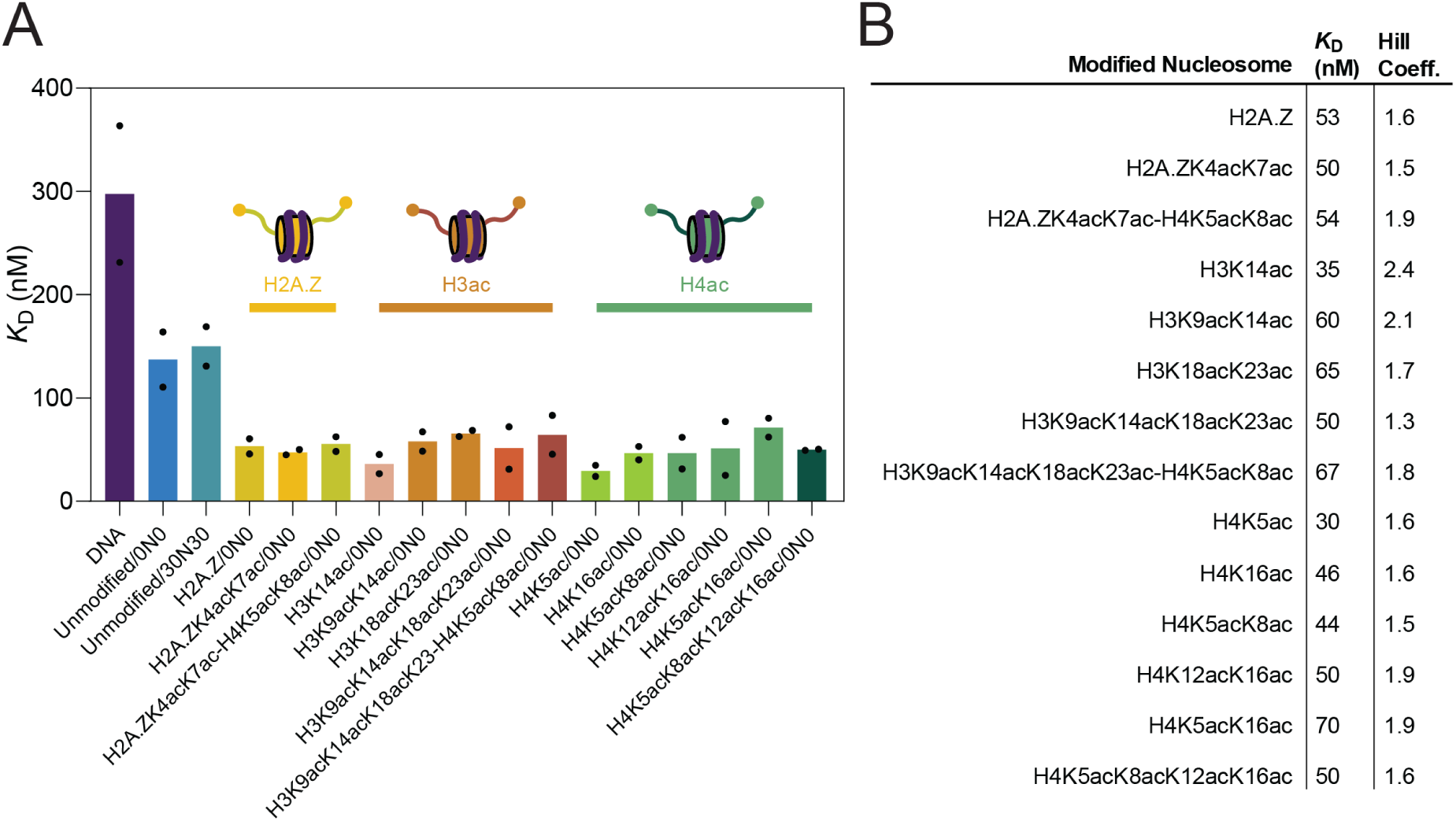
BRD4-nucleosome interactions show little preference for specific acetylation patterns. **A.** Bar plot showing mean MST affinities for BRD4S binding to nucleosomes with different modification states. Two technical replicates are shown. **B.** Table of affinities from MST assays (technical duplicates) measuring binding of BRD4S to modified nucleosomes. The SEM of all MST measurements is estimated to be 15% (see methods).

Overall, the MST and library interrogation data agree: acetylation broadly enhances the interaction between BRD4 and nucleosomes in both cases. However, the preference is significantly less than observed for interactions between isolated bromodomains and short peptides. Furthermore, there is little evidence of significant preferences for specific histone acetylation patterns.

### BRD4 makes canonical BD-AcK contacts when binding acetylated nucleosomes

Given the apparent disparity between our measurements and prior studies focusing on BD-peptide interactions, we sought to assess whether the binding mode observed in BD-peptide interactions is also observed in the context of our full-length BRD4 protein. We prepared synthetic histone H4(1–14) peptides acetylated on either K5 alone or on K5 and K8 and used surface plasmon resonance (SPR) to assess their binding to immobilized full-length BRD4S (**Figure 6A**). In close agreement with published work using isolated BRD4-BD1, we observed an affinity of 1.2 μM for BRD4S binding to H4(1–14)K5acK8ac (7 μM in previous work (9)) and a reduction of at least 50-fold in affinity for binding of H4(1–14)K5ac (*K*_D_ > 50 μM).

**Figure 6:**
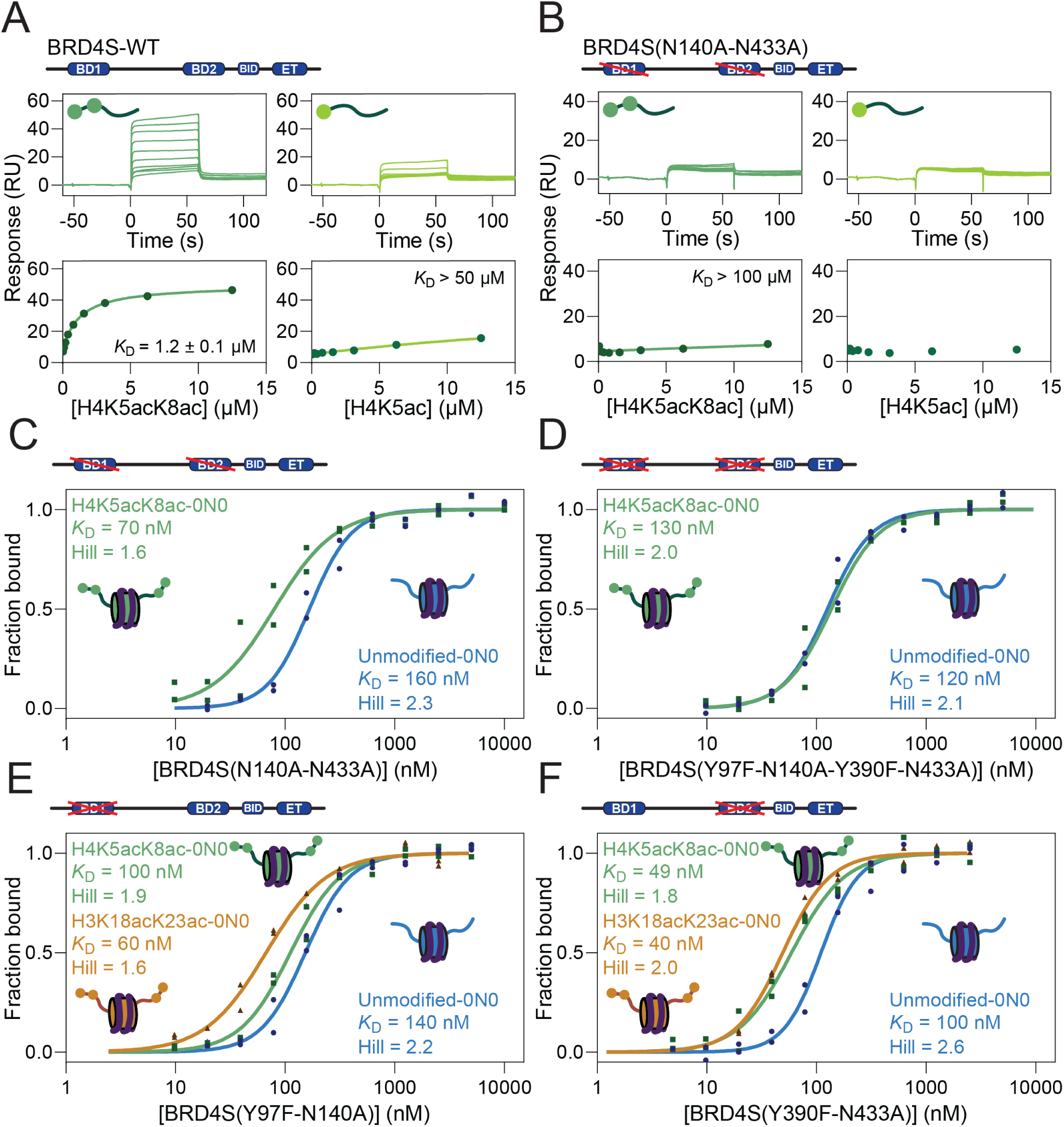
Effect of bromodomain mutation on BRD4 binding to acetylated peptides and nucleosomes. **A.** SPR data for WT BRD4S (immobilised at 3800 RU) binding to histone H4(1–14) peptides bearing either K5ac or K5acK8ac modifications. **B.** SPR data for BRD4S(N140A-N433A) (immobilised at 2500 RU) binding to H4(1–14) peptides bearing either K5ac or K5acK8ac modifications. *K*D and standard error values are calculated from a fit to a simple Langmuir 1:1 binding isotherm. **C.** MST titration of BRD4S(N140A-N433A) into unmodified nucleosomes or nucleosomes bearing either K5ac or K5acK8ac modifications. **D.** MST titration of BRD4S(Y97F-N140A-Y390F-N433A) into unmodified nucleosomes or nucleosomes bearing an H4K5acK8ac modification. **E.** MST titration of BRD4-Y97F-N140A into unmodified nucleosomes or nucleosomes bearing H3K18 and K23. **F.** MST titration of BRD4(Y390F-N433A) to unmodified nucleosomes or nucleosomes bearing either H4K5acK8ac or H3K18acK23ac modifications. The SEM of all MST measurements is estimated to be 15% (see methods).

To determine whether the preference for diacetylation was caused by canonical interactions with the AcK-binding pocket, we prepared a series of BRD4S mutants. BET bromodomains feature a conserved asparagine in the binding pocket that forms a hydrogen bond with a lysine acetyl group (**Supplementary Figure 8**) (18), and mutation of this asparagine abolishes BD binding to acetylated H4K5acK8ac peptides (42). We first prepared a double mutant [BRD4S(N140A-N433A)] in which the asparagine in both BD1 and BD2 was changed to alanine. In line with previous findings for isolated BDs, SPR titrations revealed significant decreases in the binding of H4(1–14) peptides (**Figure 6B**); H4(1–14)K5acK8ac bound at least ∼100 times weaker to the mutant than to wildtype BRD4S wild type (*K*D > 100 μM) and no binding was observed for the H4(1–14)K5ac peptide. In stark contrast, the affinity of this double mutant for either acetylated (H4K5acK8ac) or unmodified nucleosomes changed by less than two-fold compared to the wildtype protein; *K*D’s were 70 nM and 160 nM, respectively, versus 44 and 130 nM for wildtype BRD4S (**Figure 6C**).

We therefore made a second mutation in each binding pocket, targeting a conserved tyrosine that forms a water-mediated hydrogen bond with bound AcK residues (**Supplementary Figure 8**); mutation of this residue to phenylalanine also abolishes binding of BET BDs to acetylated peptides (42). MST titrations showed that this quadruple mutation [BRD4S(Y97F-N140A-Y390F-N433A)] eliminated the affinity difference between unmodified and H4K5acK8ac nucleosomes (*K*D = 120 and 130 nM, respectively) demonstrating that the two-fold preference displayed by wildtype BRD4S for acetylated nucleosomes is mediated by interactions with the AcK-binding pocket.

With the identification of mutations that disrupt acetyllysine binding in the context of a nucleosome, we next assessed the modification preferences of each BD (**Figure 6E, F**). Mutation of BD1 (Y97F-N140A) reduced the preference of BRD4S for H4K5acK8ac nucleosomes from 2-fold to 1.4-fold (*K*D = 100 nM for H4K5acK8ac and 140 nM for unmodified), whereas mutation of BD2 had no effect on this preference. No reduction in affinity for nucleosomes bearing H3K18acK23ac was observed upon mutation of either BD1 or BD2. These data indicate that some redundancy might exist in the AcK-binding properties of the two BDs, with both BD1 and BD2 able to contribute to AcK recognition in the context of full-length BRD4 binding a nucleosomal target.

Taken together, these data indicate that canonical BD-AcK interactions do take place in the context of a complex formed between full-length BRD4 and an acetylated nucleosome but suggest that these contacts make only a small contribution to binding. The data also demonstrate that the interaction of BRD4S with unmodified nucleosomes does not involve significant contacts with the AcK binding pocket.

### The BD inhibitor JQ1 does not displace BRD4 from acetylated mononucleosomes

Because the effect of BD pocket mutations was unexpectedly modest, we next assessed the impact of the small-molecule BD inhibitor JQ1 (17) on the BRD4-nucleosome interaction. JQ1, which binds competitively in the AcK-binding pocket of BET BDs, has been used in numerous *in vitro* and cell-based studies to disrupt BET protein activity (43).

SPR analysis showed that JQ1 bound immobilised full-length BRD4S with an affinity of 40 nM (consistent with published affinities for JQ1-BD interactions (17)) and gave a response at full saturation of ∼20 RU under our assay conditions (**Figure 7A**). We then tested whether JQ1 was capable of displacing H4K5acK8ac peptide from the BRD4S in a competition experiment. **Figure 7B** shows titration of JQ1 into immobilized BRD4S in the presence of a constant concentration of H4K5acK8ac peptide. The addition of JQ1 changed the response size and off-rate in line with JQ1 successfully displacing the histone H4 peptide from BRD4S.

**Figure 7:**
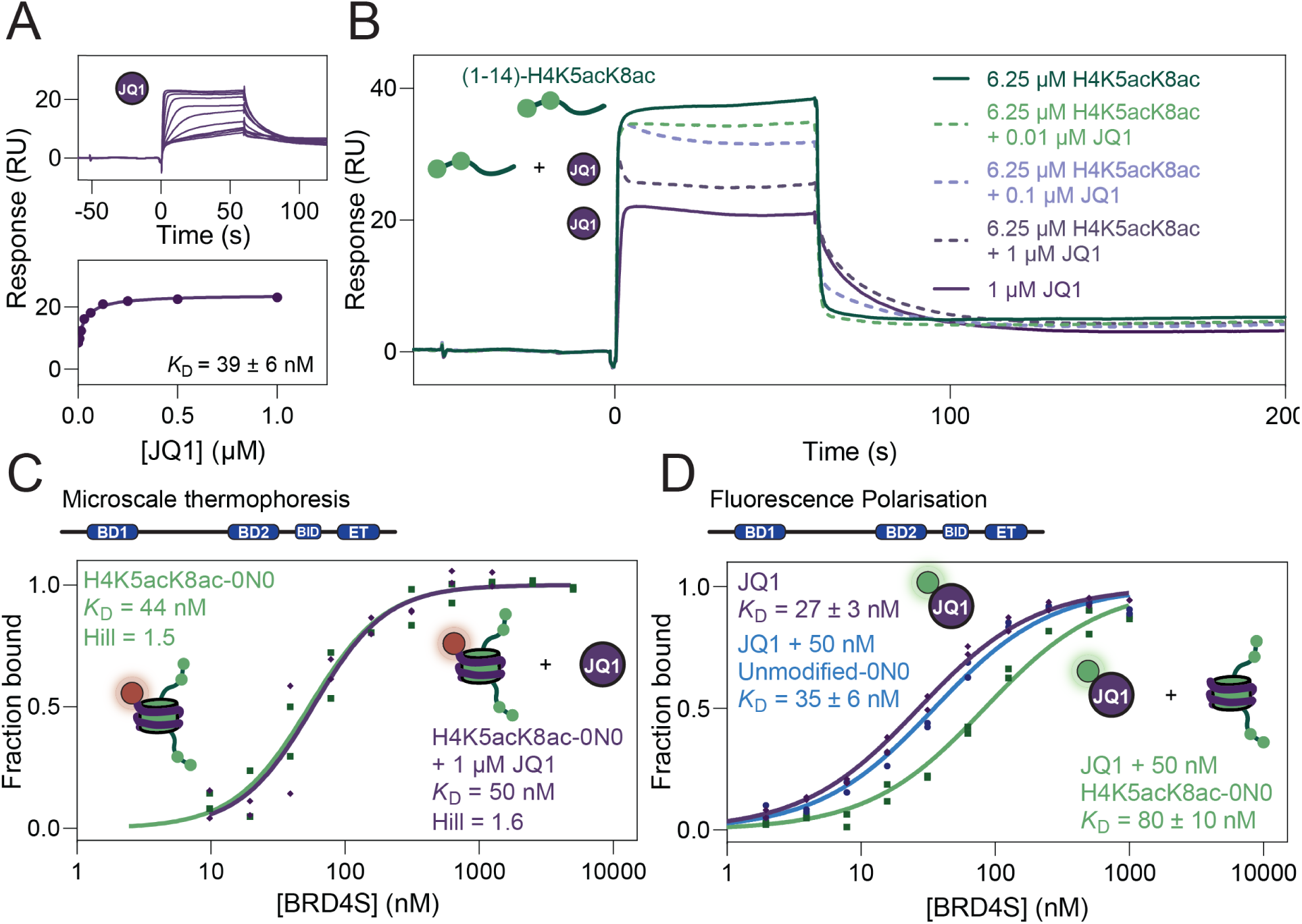
JQ1 inhibition of BRD4 binding to histone peptides and nucleosomes. **A.** SPR titration immobilized BRD4S with JQ1. *K*D and standard error values are calculated from a fit to a simple Langmuir 1:1 binding isotherm. **B.** SPR competition experiment with immobilized BRD4S, a constant concentration of H4K5acK8ac peptide (6.25 μM) and increasing concentrations of JQ1. **C.** MST titrations of BRD4S into unmodified and H4 acetylated nucleosomes, in the absence and presence of 1 μΜ JQ1. The SEM of all MST measurements is estimated to be 15% (see methods) **D.** Fluorescence polarisation titration of BRD4S on FITC-labelled JQ1 either with no nucleosome, 50 nM unmodified nucleosome or 50 nM H4K5acK8ac nucleosome. *K*D and standard error values are calculated from a fit to a simple Langmuir 1:1 binding isotherm.

In an MST competition experiment, we then titrated H4K5acK8ac nucleosomes with BRD4S in the presence of 1 μM JQ1, hypothesizing that JQ1 would block interactions with the AcK pocket, and the affinity would be reduced as it was when we examined the BRD4S quadruple mutant. Surprisingly, however, the addition of JQ1 had no measurable effect on the BRD4S-nucleosome interaction at either 75 or 150 mM NaCl (*K*D = 50 nM in 75 mM NaCl **Figure 7C, Supplementary Figure 9**). To further explore this result, we asked the converse question: whether acetylated nucleosomes could displace binding of BRD4S to JQ1. We used fluorescence polarisation to measure the affinity of FITC-labelled JQ1 for BRD4S; the measured *K*D of 27 nM (**Figure 7D**) was consistent with the affinity determined by SPR (**Figure 7A**). We then repeated the titration of BRD4S into FITC-JQ1 in the presence of either 50 nM unmodified nucleosome or 50 nM H4K5acK8ac nucleosome. The former titration yielded an essentially unchanged *K*D (35 nM), whereas the addition of acetylated nucleosome reduced the affinity by ∼3-fold (*K*D = 80 nM). These data indicate that the acetylated nucleosome could displace JQ1 from BRD4S, pointing to an asymmetry in the competition between nucleosomal H4K5acK8ac and JQ1 for the AcK pocket of BRD4 BDs.

## DISCUSSION

### BRD4-nucleosome interactions are dominated by protein-DNA contacts

To date, our understanding of BET-protein biochemistry has been built almost entirely on reductionist studies conducted using isolated domains and short acetylated peptides. To take a step closer to the cellular context, we have examined the interaction of purified full-length BRD4 with recombinant acetylated nucleosomes.

As a foundation for these experiments, we first assessed the binding of BRD4 to naked DNA – a 267-bp sequence comprising the Widom sequence with 60 bp of flanking DNA on each end. In line with two previous reports (44,45), BRD4L bound this DNA with tight affinity. Successive C-terminal truncations reduced the affinity and isolated BDs or the ET domain displayed only very weak binding, as has been observed before (46). It was notable that the BRD4S isoform bound with high apparent cooperativity; this could be related to the known ability of BRD4 to dimerize, though this would not explain the lower cooperativity shown for BRD4L.

Does this binding of BRD4 to DNA have a functional role? Rather unexpectedly, multiple reports demonstrate that the peak of the BRD4 chromatin occupancy profile at promoters corresponds precisely to the nucleosome depleted region (27,29,47–49) at which acetylation – and perhaps nucleosomes – are at low abundance or absent. BRD4 might be maintained at these sites through interactions with DNA.

It has also been suggested that non-specific BET-DNA interactions might promote association with chromatin as the proteins ‘scan’ for acetylated nucleosomes (46).

BRD4L and BRD4S bound with nanomolar affinity to recombinant unmodified nucleosomes. This finding is in line with a previous observation that used natively extracted mononucleosomes (which would have displayed a wide range of acetylation patterns (44)). The behaviour of shorter constructs, together with previous data (46,50) points to roles for the intrinsically disordered regions of BRD4 in DNA recognition.

Binding of BRD4L to nucleosomes was further enhanced by the presence of flanking DNA. A sensitivity for flanking DNA is also characteristic of several chromatin remodelling complexes (51–55). It is notable that the ET domain of BRD4 can interact with subunits from several such complexes (56,57). It is thus possible that the sensitivity of BRD4L (but not BRD4S) for flanking DNA hints at an isoform-dependent role for BRD4 in the regulation of chromatin remodelling.

### Acetylation modestly and non-selectively increases BRD4-nucleosome affinity

The impact of histone acetylation on the BRD4-nucleosome interaction was substantially weaker than we expected. In contrast to the ∼100-fold enhancements in affinity frequently observed for BD-peptide interactions, we observed increases of 2–4 fold when using 0N0 nucleosomes. Flanking DNA eliminated even this modest preference for acetylated nucleosomes. Equally striking was the lack of dependence on the pattern of acetylation. Multiple permutations of H2A.Z, H3 and/or H4 acetylation yielded similar affinity changes of 2–4-fold.

It has been previously shown that DNA and acetyllysine compete for BRD-BD1 binding, which could contribute to the modest acetylation preference that we observe (46). Transient interactions made by histone tails with the nucleosome might also play a role (58). In this context, both BPTF and the tandem PHDs of CHD4 show weaker binding to nucleosomes than to relevant histone tail peptides, and it was suggested that tail accessibility might be reduced in intact nucleosomes (59,60). In contrast, histone readers such as HP1, BPTF and PHIP show essentially no binding to unmodified nucleosomes but rather recognize only modified nucleosomes (60–62). Thus, a range of degrees of dependence exist among histone readers for their target modifications, which might point towards different functions.

How does our observed lack of specificity compare to BRD4 behaviour in cells? Perhaps surprisingly, the acetylation preferences of BRD4 *in vivo* remain unclear. ChIP-seq data in multiple studies using different cell lines show that, overall, BRD4-bound loci are enriched in acetylation and there are indications that BRD4 is more closely associated with H4K5acK8ac than with H3K27ac (10,11). However, the prominence of H4K5acK8ac in hyperacetylated chromatin makes it difficult to draw a clear conclusion and no combination of histone acetylation marks definitively predicts BRD4 chromatin occupancy (63). Other *in vitro* studies investigating the enrichment of particular modification states following pull-down of BRD4 generally support the importance of hyperacetylation in BRD4/nucleosome binding (64,65). Overall, these data indicate (a) that histone acetylation plays a role in recruitment of BRD4 to chromatin, though the enhancement in binding is more limited than other studies have suggested, and (b) that a wide range of acetylation marks might be able to play similar roles in BRD recruitment.

### The pharmacological activity of BET inhibitors might reflect the disruption of BET-transcription factor rather than BET-histone interactions

Our findings, together with published genome occupancy data, point to a model in which BRD4 binds with high affinity and low specificity to acetylated nucleosomes. This model contrasts somewhat with our current understanding of how BET proteins engage transcription factors (TFs), where affinities are likely lower (because co-binding to DNA is not involved) but sequence-specific acetylation is a critical element. For example, the interaction between BRD3-BD1 and the erythroid TF GATA1, which is required for GATA1 to find its genomic target sites and to drive erythroid differentiation, is mediated by a diacetylated KGGK motif in GATA1 (22,23). Mutation of these two lysines (but not multiple surrounding lysines that can also be acetylated) to arginine abrogates the interaction and blocks erythroid differentiation (22,24). It is notable that occupancy by GATA1 is a better predictor of BRD4 occupancy in erythroid precursor cells than histone acetylation (48), highlighting the close relationship between BRD4 and GATA1.

Closely related observations have been made for several other TFs. Diacetylation of a KGGK motif in the TF TWIST (but again not acetylation of surrounding lysines) controls its interaction with BRD4 and its cellular activity (26), and corresponding observations have been made for the myeloid TF ERG (47). This latter study also noted that essentially all BRD4-occupied sites were co-occupied by one or more TFs and highlighted the strong colocalization of BRD4 and TFs at the (nucleosome-depleted) transcriptional start site. E2F1, MyoD1, IRF1 and AR also interact with BRD4 in an acetylation dependent manner, likely through KXXK motifs (where X is any amino acid) (16,26,66–71). Finally, mutation of a single lysine (of many that can be acetylated) in the RelA subunit of the lipopolysaccharide (LPS) responsive TF NF-kB prevents its interaction with BRD4 following LPS stimulation and downregulates thousands of NF-κB-controlled genes. Together, these data indicate that site-specific, AcK-mediated interactions between TFs and BET proteins are important drivers of transcriptional regulation.

This conclusion raises a question about the mechanism of action of pharmacological BET inhibitors such as JQ1, which compete directly for the AcK binding site of BET BDs. Treatment of cells with JQ1 often leads to selective depletion of BRD4 across the genome (rather than displacement from all acetylated nucleosomes) and also leads to gene-specific changes in transcription (29,67,72), suggesting that small-molecule inhibitors disrupt some BET interactions more effectively than others.

Our data indicate that JQ1 is more effective at displacing an isolated diacetylated KXXK motif from BRD4 than it is at displacing a nucleosome bearing the same modification. We propose that binding of BRD4 binding to acetylated nucleosomes is likely multivalent due to additional contacts with DNA (perhaps mediated through intrinsically disordered regions). In such a binding mode, JQ1 is unable to displace BD-histone contacts due to the contributions of these DNA contacts compete effectively for BRD4 because of these contacts. We therefore speculate that BRD4-TF interactions might be more sensitive to pharmacological inhibition than BRD4-nucleosome interactions and therefore might represent the primary target of BET inhibitors. This idea awaits testing.

## AUTHOR CONTRIBUTIONS

Lucien S. Lambrechts and Xavier J. Reid: Conceptualisation, Formal analysis, Investigation, Methodology, Validation, Visualisation, Writing—original draft. Lucas Kambanis, Clement Luong, Erekle Kobakhidze and Yichen Zhong: Investigation, Methodology. Andrea Daners, Karishma Patel, Charlotte K. Franck, Hakimeh Moghaddas Sani and Cameron Taylor: Resources. Jason K. K. Low: Investigation, Methodology, Supervision. Richard J. Payne: Funding acquisition, Resources, Supervision. Joel P. Mackay: Conceptualisation, Funding acquisition, Project administration, Resources, Supervision, Writing—review & editing

## SUPPLEMENTARY DATA

Supplementary Data are available at NAR online.

## DATA AVAILABILITY

Data for all main figures and supplementary figure 9 are available in a source data file.

The nucleosome mass spectrometry proteomics data following BRD4 pull-down have been deposited to the ProteomeXchange Consortium via the PRIDE (41) partner repository with the dataset identifier PXD064221

## REFERENCES

1. Garner, E., Romero, P., Dunker, A.K., Brown, C. and Obradovic, Z. (1999) Predicting Binding Regions within Disordered Proteins. Genome Informatics, 10, 41–50.

2. Wang, N., Wu, R., Tang, D. and Kang, R. (2021) The BET family in immunity and disease. Signal Transduct. Target. Ther., 6, 23.

3. Donczew, R. and Hahn, S. (2021) BET family members Bdf1/2 modulate global transcription initiation and elongation in Saccharomyces cerevisiae. Elife, 10.

4. Wu, S.Y., Lee, C.F., Lai, H.T., Yu, C.T., Lee, J.E., Zuo, H., Tsai, S.Y., Tsai, M.J., Ge, K., Wan, Y. et al. (2020) Opposing Functions of BRD4 Isoforms in Breast Cancer. Mol. Cell, 78, 1114–1132.e1110.

5. Patel, M.C., Debrosse, M., Smith, M., Dey, A., Huynh, W., Sarai, N., Heightman, T.D., Tamura, T. and Ozato, K. (2013) BRD4 coordinates recruitment of pause release factor P-TEFb and the pausing complex NELF/DSIF to regulate transcription elongation of interferon-stimulated genes. Mol. Cell. Biol., 33, 2497–2507.

6. Wang, R., Li, Q., Helfer, C.M., Jiao, J. and You, J. (2012) Bromodomain Protein Brd4 Associated with Acetylated Chromatin Is Important for Maintenance of Higher-order Chromatin Structure. J. Biol. Chem., 287, 10738–10752.

7. Shang, E., Wang, X., Wen, D., Greenberg, D.A. and Wolgemuth, D.J. (2009) Double bromodomain-containing gene Brd2 is essential for embryonic development in mouse. Dev. Dyn., 238, 908–917.

8. Umehara, T., Nakamura, Y., Jang, M.K., Nakano, K., Tanaka, A., Ozato, K., Padmanabhan, B. and Yokoyama, S. (2010) Structural basis for acetylated histone H4 recognition by the human BRD2 bromodomain. J. Biol. Chem., 285, 7610–7618.

9. Filippakopoulos, P., Picaud, S., Mangos, M., Keates, T., Lambert, J.-P., Barsyte-Lovejoy, D., Felletar, I., Volkmer, R., Müller, S., Pawson, T. et al. (2012) Histone Recognition and Large-Scale Structural Analysis of the Human Bromodomain Family. Cell, 149, 214–231.

10. Liu, W., Ma, Q., Wong, K., Li, W., Ohgi, K., Zhang, J., Aneel and Michael. (2013) Brd4 and JMJD6-Associated Anti-Pause Enhancers in Regulation of Transcriptional Pause Release. Cell, 155, 1581–1595.

11. Handoko, L., Kaczkowski, B., Hon, C.-C., Lizio, M., Wakamori, M., Matsuda, T., Ito, T., Jeyamohan, P., Sato, Y., Sakamoto, K. et al. (2018) JQ1 affects BRD2-dependent and independent transcription regulation without disrupting H4-hyperacetylated chromatin states. Epigenetics, 13, 410–431.

12. Sanchez, R., Meslamani, J. and Zhou, M.-M. (2014) The bromodomain: From epigenome reader to druggable target. Biochim. Biophys. Acta, 1839, 676–685.

13. Lambert, J.-P., Picaud, S., Fujisawa, T., Hou, H., Savitsky, P., Uusküla-Reimand, L., Gupta, G.D., Abdouni, H., Lin, Z.-Y., Tucholska, M. et al. (2019) Interactome Rewiring Following Pharmacological Targeting of BET Bromodomains. Mol. Cell, 73, 621–638.e617.

14. Huang, B., Yang, X.-D., Zhou, M.-M., Ozato, K. and Chen, L.-F. (2009) Brd4 Coactivates Transcriptional Activation of NF-κB via Specific Binding to Acetylated RelA. Mol. Cell. Biol., 29, 1375–1387.

15. Gamsjaeger, R., Webb, S.R., Lamonica, J.M., Billin, A., Blobel, G.A. and Mackay, J.P. (2011) Structural Basis and Specificity of Acetylated Transcription Factor GATA1 Recognition by BET Family Bromodomain Protein Brd3. Mol. Cell. Biol., 31, 2632–2640.

16. Patel, K., Solomon, P.D., Walshe, J.L., Ford, D.J., Wilkinson-White, L., Payne, R.J., Low, J.K.K. and Mackay, J.P. (2021) BET-Family Bromodomains Can Recognize Diacetylated Sequences from Transcription Factors Using a Conserved Mechanism. Biochemistry, 60, 648–662.

17. Filippakopoulos, P., Qi, J., Picaud, S., Shen, Y., Smith, W.B., Fedorov, O., Morse, E.M., Keates, T., Hickman, T.T., Felletar, I. et al. (2010) Selective inhibition of BET bromodomains. Nature, 468, 1067–1073.

18. Morinière, J., Rousseaux, S., Steuerwald, U., Soler-López, M., Curtet, S., Vitte, A.-L., Govin, J., Gaucher, J., Sadoul, K., Hart, D.J. et al. (2009) Cooperative binding of two acetylation marks on a histone tail by a single bromodomain. Nature, 461, 664–668.

19. Patel, K., Solomon, P.D., Walshe, J.L., Low, J.K.K. and Mackay, J.P. (2021) The bromodomains of BET family proteins can recognize diacetylated histone H2A.Z. Protein Sci., 30, 464–476.

20. Huang, B., Yang, X.D., Zhou, M.M., Ozato, K. and Chen, L.F. (2009) Brd4 coactivates transcriptional activation of NF-kappaB via specific binding to acetylated RelA. Mol. Cell. Biol., 29, 1375–1387.

21. Wollebo, H.S., Bellizzi, A., Cossari, D.H., Salkind, J., Safak, M. and White, M.K. (2016) The Brd4 acetyllysine-binding protein is involved in activation of polyomavirus JC. J. Neurovirol., 22, 615–625.

22. Lamonica, J.M., Deng, W., Kadauke, S., Campbell, A.E., Gamsjaeger, R., Wang, H., Cheng, Y., Billin, A.N., Hardison, R.C., Mackay, J.P. et al. (2011) Bromodomain protein Brd3 associates with acetylated GATA1 to promote its chromatin occupancy at erythroid target genes. Proc. Natl. Acad. Sci. U. S. A., 108, E159–E168.

23. Stonestrom, A.J., Hsu, S.C., Jahn, K.S., Huang, P., Keller, C.A., Giardine, B.M., Kadauke, S., Campbell, A.E., Evans, P., Hardison, R.C. et al. (2015) Functions of BET proteins in erythroid gene expression. Blood, 125, 2825–2834.

24. Hung, H.-L., Lau, J., Kim, A.Y., Weiss, M.J. and Blobel, G.A. (1999) CREB-Binding Protein Acetylates Hematopoietic Transcription Factor GATA-1 at Functionally Important Sites. Mol. Cell. Biol., 19, 3496–3505.

25. Zou, Z., Huang, B., Wu, X., Zhang, H., Qi, J., Bradner, J., Nair, S. and Chen, L.F. (2014) Brd4 maintains constitutively active NF-κB in cancer cells by binding to acetylated RelA. Oncogene, 33, 2395–2404.

26. Shi, J., Wang, Y., Zeng, L., Wu, Y., Deng, J., Zhang, Q., Lin, Y., Li, J., Kang, T., Tao, M. et al. (2014) Disrupting the Interaction of BRD4 with Diacetylated Twist Suppresses Tumorigenesis in Basal-like Breast Cancer. Cancer Cell, 25, 210–225.

27. Nicodeme, E., Jeffrey, K.L., Schaefer, U., Beinke, S., Dewell, S., Chung, C.-W., Chandwani, R., Marazzi, I., Wilson, P., Coste, H. et al. (2010) Suppression of inflammation by a synthetic histone mimic. Nature, 468, 1119–1123.

28. Dawson, M.A., Prinjha, R.K., Dittmann, A., Giotopoulos, G., Bantscheff, M., Chan, W.I., Robson, S.C., Chung, C.W., Hopf, C., Savitski, M.M. et al. (2011) Inhibition of BET recruitment to chromatin as an effective treatment for MLL-fusion leukaemia. Nature, 478, 529–533.

29. Lovén, J., Heather, Charles, Lau, A., David, Christopher, James, Tong and Richard. (2013) Selective Inhibition of Tumor Oncogenes by Disruption of Super-Enhancers. Cell, 153, 320–334.

30. Wang, Z.-Q., Zhang, Z.-C., Wu, Y.-Y., Pi, Y.-N., Lou, S.-H., Liu, T.-B., Lou, G. and Yang, C. (2023) Bromodomain and extraterminal (BET) proteins: biological functions, diseases and targeted therapy. Signal Transduct. Target. Ther., 8.

31. Zhong, Y., Paudel, B.P., Ryan, D.P., Low, J.K.K., Franck, C., Patel, K., Bedward, M.J., Torrado, M., Payne, R.J., Van Oijen, A.M., et al. (2020) CHD4 slides nucleosomes by decoupling entry- and exit-side DNA translocation. Nat. Commun., 11.

32. Studier, F.W. (2005) Protein production by auto-induction in high density shaking cultures. Protein Expr. Purif., 41, 207–234.

33. Miller, G.M., Flynn, E.M., Tom, J., Song, A. and Cochran, A.G. (2022) Trifluoroacetyl Lysine as a Bromodomain Binding Mimic of Lysine Acetylation. ACS Chem. Biol., 17, 1022–1029.

34. Franck, C., Foster, S.R., Johansen-Leete, J., Chowdhury, S., Cielesh, M., Bhusal, R.P., Mackay, J.P., Larance, M., Stone, M.J. and Payne, R.J. (2020) Semisynthesis of an evasin from tick saliva reveals a critical role of tyrosine sulfation for chemokine binding and inhibition. Proc. Natl. Acad. Sci. U. S. A., 117, 12657–12664.

35. Thåström, A., Lowary, P.T., Widlund, H.R., Cao, H., Kubista, M. and Widom, J. (1999) Sequence motifs and free energies of selected natural and non-natural nucleosome positioning DNA sequences11Edited by T. Richmond. J. Mol. Biol., 288, 213–229.

36. Patel, K., Walport, L.J., Walshe, J.L., Solomon, P.D., Low, J.K.K., Tran, D.H., Mouradian, K.S., Silva, A.P.G., Wilkinson-White, L., Norman, A. et al. (2020) Cyclic peptides can engage a single binding pocket through highly divergent modes. Proc. Natl. Acad. Sci. U. S. A., 117, 26728–26738.

37. Hughes, C.S., Foehr, S., Garfield, D.A., Furlong, E.E., Steinmetz, L.M. and Krijgsveld, J. (2014) Ultrasensitive proteome analysis using paramagnetic bead technology. Mol. Syst. Biol., 10, 757.

38. MacLean, B., Tomazela, D.M., Shulman, N., Chambers, M., Finney, G.L., Frewen, B., Kern, R., Tabb, D.L., Liebler, D.C. and MacCoss, M.J. (2010) Skyline: an open source document editor for creating and analyzing targeted proteomics experiments. Bioinformatics, 26, 966–968.

39. Frewen, B. and MacCoss, M.J. (2007) Using BiblioSpec for creating and searching tandem MS peptide libraries. Curr. Protoc. Bioinform., Chapter 13, 13 17 11–13 17 12.

40. Lin, S. and Garcia, B.A. (2012) Examining histone posttranslational modification patterns by high-resolution mass spectrometry. Methods Enzymol., 512, 3–28.

41. Perez-Riverol, Y., Bandla, C., Kundu, Deepti J., Kamatchinathan, S., Bai, J., Hewapathirana, S., John, Nithu S., Prakash, A., Walzer, M., Wang, S. et al. (2025) The PRIDE database at 20 years: 2025 update. Nucleic Acids Res., 53, D543–D553.

42. Jung, M., Philpott, M., Müller, S., Schulze, J., Badock, V., Eberspächer, U., Moosmayer, D., Bader, B., Schmees, N., Fernández-Montalván, A. et al. (2014) Affinity Map of Bromodomain Protein 4 (BRD4) Interactions with the Histone H4 Tail and the Small Molecule Inhibitor JQ1*. J. Biol. Chem., 289, 9304–9319.

43. Garnier, J.-M., Sharp, P.P. and Burns, C.J. (2014) BET bromodomain inhibitors: a patent review. Expert Opin. Ther. Pat., 24, 185–199.

44. Larue, R.C., Plumb, M.R., Crowe, B.L., Shkriabai, N., Sharma, A., Difiore, J., Malani, N., Aiyer, S.S., Roth, M.J., Bushman, F.D. et al. (2014) Bimodal high-affinity association of Brd4 with murine leukemia virus integrase and mononucleosomes. Nucleic Acids Res., 42, 4868–4881.

45. Han, X., Yu, D., Gu, R., Jia, Y., Wang, Q., Jaganathan, A., Yang, X., Yu, M., Babault, N., Zhao, C. et al. (2020) Roles of the BRD4 short isoform in phase separation and active gene transcription. Nat. Struct. Mol. Biol., 27, 333–341.

46. Kalra, P., Zahid, H., Ayoub, A., Dou, Y. and Pomerantz, W.C.K. (2022) Alternative Mechanisms for DNA Engagement by BET Bromodomain-Containing Proteins. Biochemistry, 61, 1260–1272.

47. Roe, J.-S., Mercan, F., Rivera, K., Darryl and Christopher. (2015) BET Bromodomain Inhibition Suppresses the Function of Hematopoietic Transcription Factors in Acute Myeloid Leukemia. Mol. Cell, 58, 1028–1039.

48. Behera, V., Stonestrom, A.J., Hamagami, N., Hsiung, C.C., Keller, C.A., Giardine, B., Sidoli, S., Yuan, Z.-F., Bhanu, N.V., Werner, M.T. et al. (2019) Interrogating Histone Acetylation and BRD4 as Mitotic Bookmarks of Transcription. Cell Rep., 27, 400–415.e405.

49. Brown, Jonathan D., Lin, Charles Y., Duan, Q., Griffin, G., Federation, A.J., Paranal, Ronald M., Bair, S., Newton, G., Lichtman, A.H., Kung, A.L. et al. (2014) NF-κB Directs Dynamic Super Enhancer Formation in Inflammation and Atherogenesis. Mol. Cell, 56, 219–231.

50. Miller, T.C.R., Simon, B., Rybin, V., Grötsch, H., Curtet, S., Khochbin, S., Carlomagno, T. and Müller, C.W. (2016) A bromodomain–DNA interaction facilitates acetylation-dependent bivalent nucleosome recognition by the BET protein BRDT. Nat. Commun., 7, 13855.

51. Leonard, John D. and Narlikar, Geeta J. (2015) A Nucleotide-Driven Switch Regulates Flanking DNA Length Sensing by a Dimeric Chromatin Remodeler. Mol. Cell, 57, 850–859.

52. Yang, J.G., Madrid, T.S., Sevastopoulos, E. and Narlikar, G.J. (2006) The chromatin-remodeling enzyme ACF is an ATP-dependent DNA length sensor that regulates nucleosome spacing. Nat. Struct. Mol. Biol., 13, 1078–1083.

53. McKnight, J.N., Jenkins, K.R., Nodelman, I.M., Escobar, T. and Bowman, G.D. (2011) Extranucleosomal DNA binding directs nucleosome sliding by Chd1. Mol. Cell. Biol., 31, 4746–4759.

54. Ocampo, J., Chereji, R.V., Eriksson, P.R. and Clark, D.J. (2016) The ISW1 and CHD1 ATP-dependent chromatin remodelers compete to set nucleosome spacing in vivo. Nucleic Acids Res., 44, 4625–4635.

55. Ocampo, J., Chereji, R.V., Eriksson, P.R. and Clark, D.J. (2019) Contrasting roles of the RSC and ISW1/CHD1 chromatin remodelers in RNA polymerase II elongation and termination. Genome Res., 29, 407–417.

56. Wai, D.C.C., Szyszka, T.N., Campbell, A.E., Kwong, C., Wilkinson-White, L.E., Silva, A.P.G., Low, J.K.K., Kwan, A.H., Gamsjaeger, R., Chalmers, J.D. et al. (2018) The BRD3 ET domain recognizes a short peptide motif through a mechanism that is conserved across chromatin remodelers and transcriptional regulators. J. Biol. Chem., 293, 7160–7175.

57. Rahman, S., Sowa, M.E., Ottinger, M., Smith, J.A., Shi, Y., Harper, J.W. and Howley, P.M. (2011) The Brd4 Extraterminal Domain Confers Transcription Activation Independent of pTEFb by Recruiting Multiple Proteins, Including NSD3. Mol. Cell. Biol., 31, 2641–2652.

58. Furukawa, A., Wakamori, M., Arimura, Y., Ohtomo, H., Tsunaka, Y., Kurumizaka, H., Umehara, T. and Nishimura, Y. (2020) Acetylated histone H4 tail enhances histone H3 tail acetylation by altering their mutual dynamics in the nucleosome. Proc. Natl. Acad. Sci. U. S. A., 117, 19661–19663.

59. Gatchalian, J., Wang, X., Ikebe, J., Cox, K.L., Tencer, A.H., Zhang, Y., Burge, N.L., Di, L., Gibson, M.D., Musselman, C.A. et al. (2017) Accessibility of the histone H3 tail in the nucleosome for binding of paired readers. Nat. Commun., 8, 1489.

60. Marunde, M.R., Fuchs, H.A., Burg, J.M., Popova, I.K., Vaidya, A., Hall, N.W., Weinzapfel, E.N., Meiners, M.J., Watson, R., Gillespie, Z.B. et al. (2024) Nucleosome conformation dictates the histone code. Elife, 13.

61. Munari, F., Soeroes, S., Zenn, H.M., Schomburg, A., Kost, N., Schröder, S., Klingberg, R., Rezaei-Ghaleh, N., Stützer, A., Gelato, K.A. et al. (2012) Methylation of Lysine 9 in Histone H3 Directs Alternative Modes of Highly Dynamic Interaction of Heterochromatin Protein hHP1β with the Nucleosome. J. Biol. Chem., 287, 33756–33765.

62. Morgan, M.A.J., Popova, I.K., Vaidya, A., Burg, J.M., Marunde, M.R., Rendleman, E.J., Dumar, Z.J., Watson, R., Meiners, M.J., Howard, S.A. et al. (2021) A trivalent nucleosome interaction by PHIP/BRWD2 is disrupted in neurodevelopmental disorders and cancer. Genes Dev., 35, 1642–1656.

63. Zippo, A., Serafini, R., Rocchigiani, M., Pennacchini, S., Krepelova, A. and Oliviero, S. (2009) Histone Crosstalk between H3S10ph and H4K16ac Generates a Histone Code that Mediates Transcription Elongation. Cell, 138, 1122–1136.

64. Nguyen, U.T.T., Bittova, L., Müller, M.M., Fierz, B., David, Y., Houck-Loomis, B., Feng, V., Dann, G.P. and Muir, T.W. (2014) Accelerated chromatin biochemistry using DNA-barcoded nucleosome libraries. Nat. Methods, 11, 834–840.

65. Lukauskas, S., Tvardovskiy, A., Nguyen, N.V., Stadler, M., Faull, P., Ravnsborg, T., Özdemir Aygenli, B., Dornauer, S., Flynn, H., Lindeboom, R.G.H. et al. (2024) Decoding chromatin states by proteomic profiling of nucleosome readers. Nature, 627, 671–679.

66. Duquet, A., Polesskaya, A., Cuvellier, S., Ait-Si-Ali, S., Héry, P., Pritchard, L.L., Gerard, M. and Harel-Bellan, A. (2006) Acetylation is important for MyoD function in adult mice. EMBO Rep., 7, 1140–1146.

67. Winter, G.E., Mayer, A., Buckley, D.L., Erb, M.A., Roderick, J.E., Vittori, S., Reyes, J.M., Di Iulio, J., Souza, A., Ott, C.J., et al. (2017) BET Bromodomain Proteins Function as Master Transcription Elongation Factors Independent of CDK9 Recruitment. Mol. Cell, 67, 5–18.e19.

68. Martínez-Balbás, M.A., Bauer, U.-M., Nielsen, S.J., Brehm, A. and Kouzarides, T. (2000) Regulation of E2F1 activity by acetylation. EMBO J., 19, 662–671.

69. Xiong, Y., Li, L., Zhang, L., Cui, Y., Wu, C., Li, H., Chen, K., Yang, Q., Xiang, R., Hu, Y. et al. (2019) The bromodomain protein BRD4 positively regulates necroptosis via modulating MLKL expression. Cell Death Differ., 26, 1929–1941.

70. Asangani, I.A., Dommeti, V.L., Wang, X., Malik, R., Cieslik, M., Yang, R., Escara-Wilke, J., Wilder-Romans, K., Dhanireddy, S., Engelke, C. et al. (2014) Therapeutic targeting of BET bromodomain proteins in castration-resistant prostate cancer. Nature, 510, 278–282.

71. Faivre, E.J., McDaniel, K.F., Albert, D.H., Mantena, S.R., Plotnik, J.P., Wilcox, D., Zhang, L., Bui, M.H., Sheppard, G.S., Wang, L. et al. (2020) Selective inhibition of the BD2 bromodomain of BET proteins in prostate cancer. Nature, 578, 306–310.

72. Shi, J. and Vakoc, C. (2014) The Mechanisms behind the Therapeutic Activity of BET Bromodomain Inhibition. Mol. Cell, 54, 728–736.

