## Supplementary Information for "The BRD4-nucleosome interaction is enhanced modestly and non-selectively by histone acetylation"

**SUPPLEMENTARY DATA**

0w0

Cy5

5’CTGCAGAAGCTTGGTCCCGGGGCCGCTCAATTGGTCGTAGCAAGCTCTAGATCCGCTTAATCGAAC

GTACGCGcTGTCCCCCGCGTTTTAACCGCCAAGGGGATTACTCCCTAGTCTCCAGGCACGTGTCAGAT

ATATACATCCTGT 3’

30w30

Cy5

5’CTCGGTACCCGGACCCTATACGCGGGCGCACTGCAGAAGCTTGGTCCCGGGGCCGCTCAATTGGTC

GTAGCAAGCTCTAGATCCGCTTAATCGAACGTACGCGcTGTCCCCCGCGTTTTAACCGCCAAGGGGAT

TACTCCCTAGTCTCCAGGCACGTGTCAGATATATACATCCTGTGCATGTAGGGGATTCTCTAGAGTCG

ACCTG 3’

46w60

Cy5

5’ATAGGGCGAATTCGAGCTCGGTACCCGGACCCTATACGCGGGCGCACTGCAGAAGCTTGGTCCCGG

GGCCGCTCAATTGGTCGTAGCAAGCTCTAGATCCGCTTAATCGAACGTACGCGcTGTCCCCCGCGTTT

TAACCGCCAAGGGGATTACTCCCTAGTCTCCAGGCACGTGTCAGATATATACATCCTGTGCATGTAGG

GGATTCTCTAGAGTCGACCTGCAGGCATGCAAGCTTGAGTATTCTATAGT 3’

60w60

Cy5

5’TAATACGACTCACTATAGGGCGAATTCGAGCTCGGTACCCGGACCCTATACGCGGGCGCACTGCAG

AAGCTTGGTCCCGGGGCCGCTCAATTGGTCGTAGCAAGCTCTAGATCCGCTTAATCGAACGTACGCGc

TGTCCCCCGCGTTTTAACCGCCAAGGGGATTACTCCCTAGTCTCCAGGCACGTGTCAGATATATACAT

CCTGTGCATGTAGGGGATTCTCTAGAGTCGACCTGCAGGCATGCAAGCTTGAGTATTCTATAGT 3’

**Supplementary Figure 1: Sequences of oligos used.** All oligos were prepared by PCR using a 5’ Cy5 labelled primer purchased from Integrated DNA Technologies. The core 601 sequence is highlighted in grey.

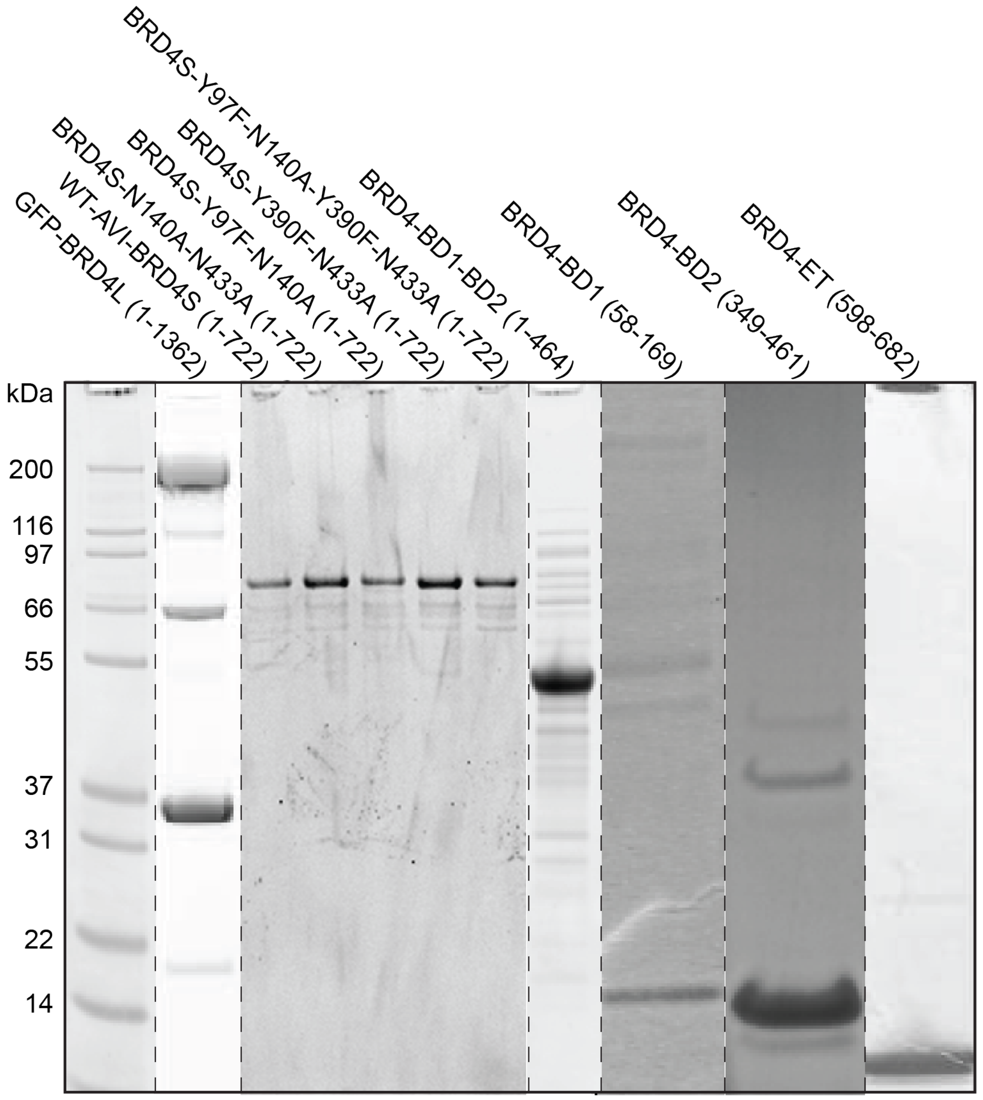


**Supplementary Figure 2: Assessment of purity of BRD4 isoforms, truncations and mutations.** A GFP-BRD4L pcDNA6 construct was expressed in HEK Expi293 cells and purified with streptavidin beads pre-incubated with Strep-GFP nanobody followed by elution with biotin. GST-3C-AVI-BRD4S pQE80L constructs were expressed in Rosetta 2(DE3) *Escherichia coli* cells and purified with GSH-Sepharose beads followed by cleavage and elution with HRV-3C protease. Proteins were purified with an additional cation exchange protocol. GST-3C-AVI-BRD4-BD1, GST-3C-AVI-BRD4-BD2, GST-3C-AVI-BRD4-BD1-BD2 fusion genes encoded in pQE80L were expressed in *Escherichia coli* BL21(DE3) cells. All constructs were purified with GSH-Sepharose beads followed by cleavage and elution with HRV-3C protease. Proteins were purified with an additional size exclusion protocol. HA-GST-3C-BRD4-ET fusion gene encoded in pGEX was expressed in *Escherichia coli* BL21(DE3) cells and were purified with GSH-Sepharose beads followed by cleavage and elution with HRV-3C protease. Representative gels are shown.

| | | |
|---|---|---|
| 1 | MSAESGPGTRLRNLPVMGDGLETSQMSTTQAQAQPQPANAASTNPPPPETSNPNKPKRQT | 60 |
| 61 | NQLQYLLRVVLKTLWKHQFAWPFQQPVDAVKLNLPDYYKIIKTPMDMGTIKKRLENNYYW | 120 |
| 121 | NAQECIQDFNTMFTNCYIYNKPGDDIVLMAEALEKLFLQKINELPTEETEIMIVQAKGRG | 180 |
| 181 | RGRKETGTAKPGVSTVPNTTQASTPPQTQTPQPNPPPVQATPHPFPAVTPDLIVQTPVMT | 240 |
| 241 | VVPPQPLQTPPPVPPQPQPPPAPAPQPVQSHPPIIAATPQPVKTKKGVKRKADTTTPTTI | 300 |
| 301 | DPIHEPPSLPPEPKTTKLGQRRESSRPVKPPKKDVPDSQQHPAPEKSSKVSEQLKCCSGI | 360 |
| 361 | LKEMFAKKHAAYAWPFYKPVDVEALGLHDYCDIIKHPMDMSTIKSKLEAREYRDAQEFGA | 420 |
| 421 | DVRLMFSNCYKYNPPDHEVVAMARKLQDVFEMRFAKMPDEPEEPVVAVSSPAVPPPTKVV | 480 |
| 481 | APPSSSDSSSDSSSDSDSSTDDSEEERAQRLAELQEQLKAVHEQLAALSQPQQNKPKKKE | 540 |
| 541 | KDKKEKKKEKHKRKEEVEENKKSKAKEPPPKKTKKNNSSNSNVSKKEPAPMKSKPPPTYE | 600 |
| 601 | SEEEDKCKPMSYEEKRQLSLDINKLPGEKLGRVVHIIQSREPSLKNSNPDEIEIDFETLK | 660 |
| 661 | PSTLRELERYVTSCLRKKRKPQAEKVDVIAGSSKMKGFSSSESESSSESSSSDSEDSETE | 720 |
| 721 | MAPKSKKKGHPGREQKKHHHHHQQMQQAPAPVPQQPPPPPQQPPPPPPPQQQQQPPPPP | 780 |
| 781 | PPPSMPQQAAPAMKSSPPPFIATQVPVLEPQLPGSVFDPIGHFTQPILHLPQPELPPHLP | 840 |
| 841 | QPPEHSTPPHLNQHAVVSPPALHNALPQQPSRPSNRAAALPPKPARPPAVSPALTQTPLL | 900 |
| 901 | PQPPMAQPPQVLLEDEEPPAPPLTSMQMQLYLQQLQKVQPPTPLLPSVKVQSQPPPPLPP | 960 |
| 961 | PPHPSVQQQLQQQPPPPPPPQPQPPPQQQHQPPPRPVHLQPMQFSTHIQQPPPPQGQQPP | 1020 |
| 1021 | HPPPGQQPPPPQPAKPQQVIQHHHSPRHHKSDPYSTGHLREAPSPLMIHSPQMSQFQSLT | 1080 |
| 1081 | HQSPPQQNVQPKKQELRAASVVQPQPLVVVKEEKIHSPIIRSEPFSPSLRPEPPKHPESI | 1140 |
| 1141 | KAPVHLPQRPEMKPVDVGRPVIRPPEQNAPPPGAPDKDKQKQEPKTPVAPKKDLKIKNMG | 1200 |
| 1201 | SWASLVQKHPTTPSSTAKSSSDSFEQFRRAAREKEEREKALKAQAEHAEKEKERLRQERM | 1260 |
| 1261 | RSREDEDALEQARRAHEEARRRQEQQQQRQEQQQQQQQAAAVAAAATPQAQSSQPQSM | 1320 |
| 1321 | LDQQRELARKREQERRRREAMAATIDMNFQSDLLSIFEENLF | 1362 |

**Supplementary Figure 3: Sequence of BRD4L with charged residues highlighted.** Positively charged residues (arginine and lysine) in BRD4L linker regions are highlighted in blue and negative residues (glutamate and aspartate) in red. Domains (BD1, BD2, ET) and predicted ordered regions (BID, CTD) are highlighted in grey.

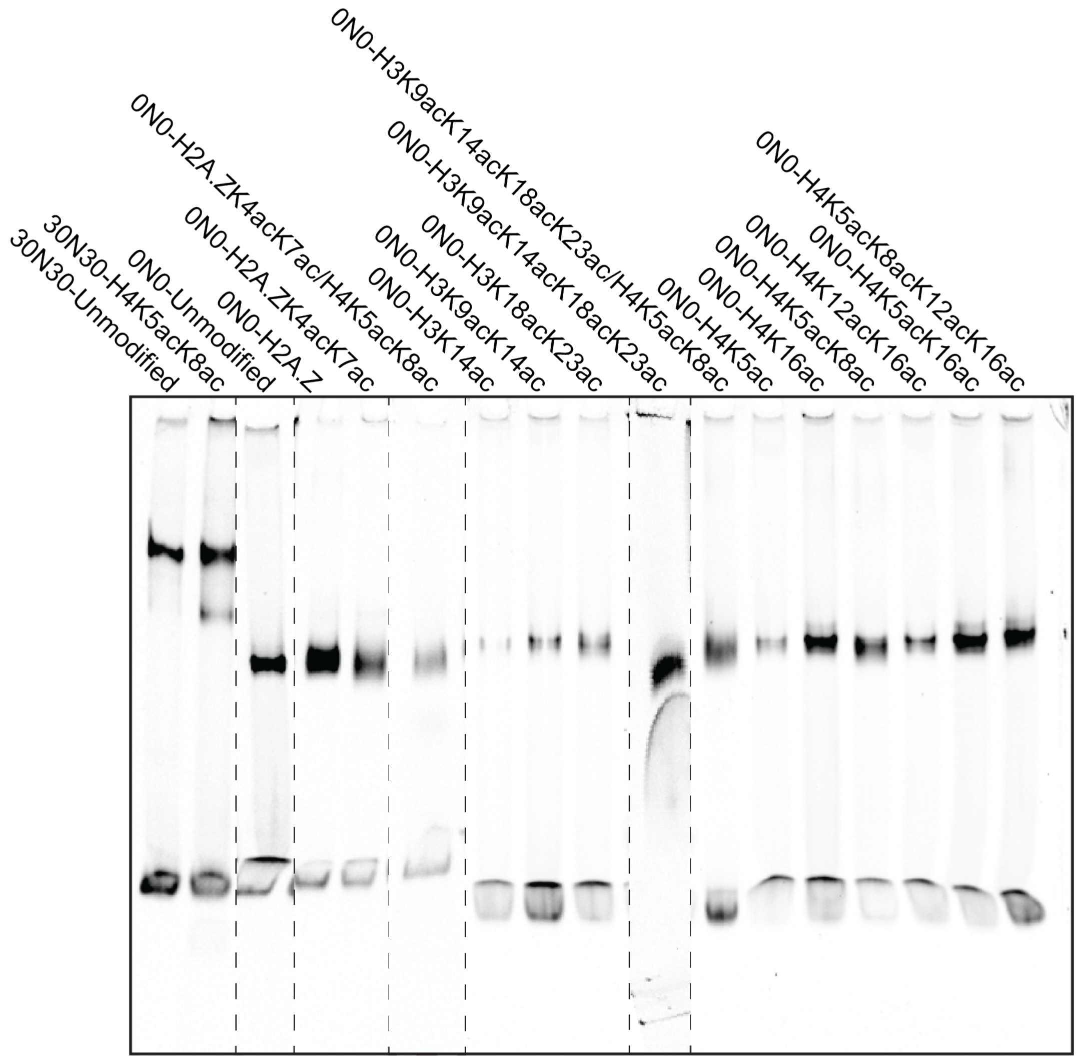


**Supplementary Figure 4: Assessment of quality of nucleosome reconstitution.** Nucleosomes were prepared by mixing histone octamer and Cy5-labelled DNA in 2 M NaCl and reconstituted using a salt-gradient dialysis protocol. Reconstituted nucleosomes were centrifuged and loaded onto 0.5× TBE 7.5% polyacrylamide gels. Gels were imaged on a Typhoon FLA-9000 laser scanner.

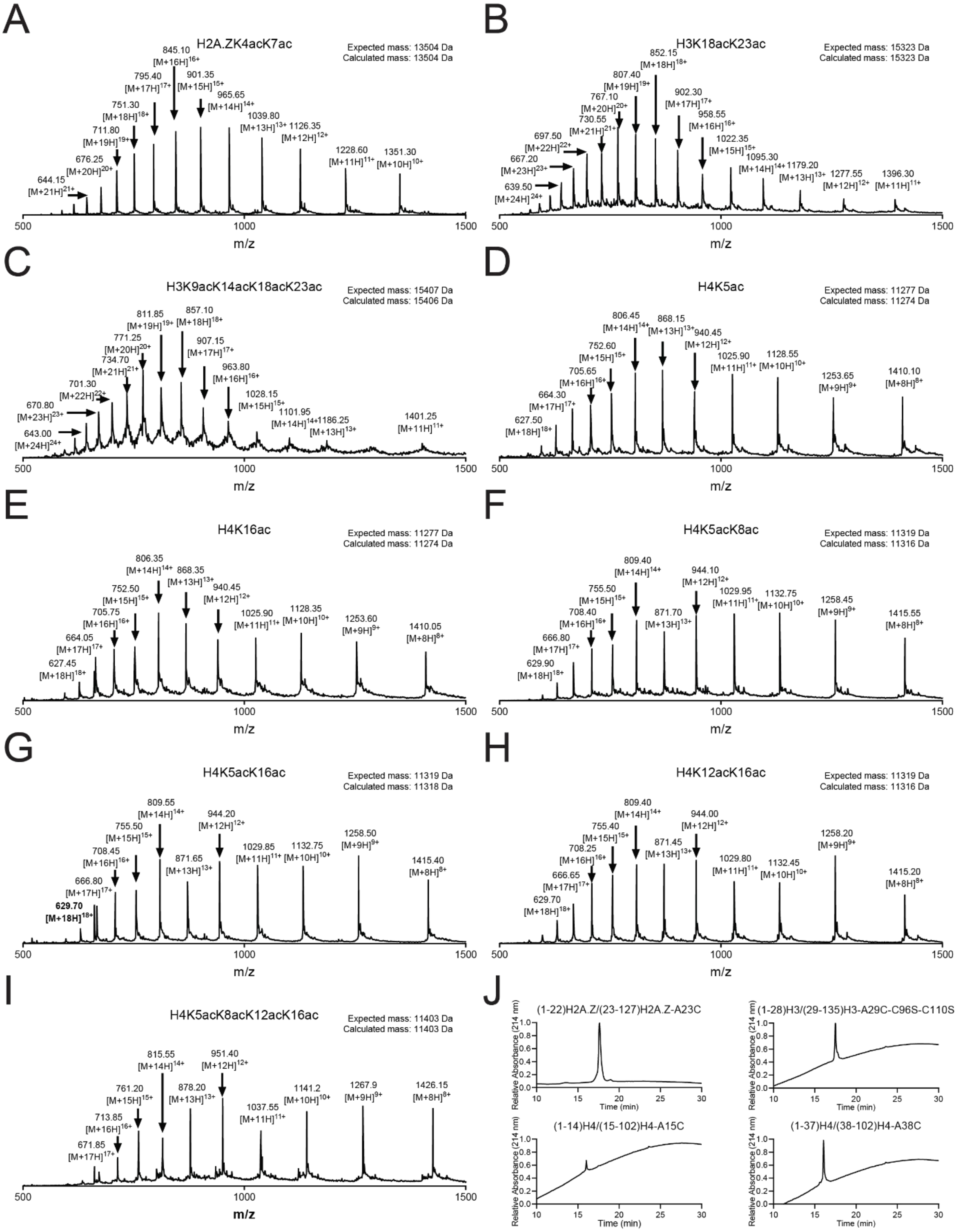


**Supplementary Figure 5: ESI-MS traces for semi-synthetic histones and analytical HPLC traces for each NCL junction. A.** ESI-MS trace for H2A.ZK4acK7ac, Calculated mass is within 3 Da of expected. **B.** ESI-MS trace for H3K18acK23ac, Calculated mass is within 3 Da of expected. **C.** ESI-MS trace for H3K9acK14acK18acK23ac, Calculated mass is within 3 Da of expected. **D.** ESI-MS trace for H4K5ac, Calculated mass is within 3 Da of expected. **E.** ESI-MS trace for H4K16ac, Calculated mass is within 3 Da of expected. **F.** ESI-MS trace for H4K5acK8ac, Calculated mass is within 3 Da of expected. **G.** ESI-MS trace for H4K5acK16ac, Calculated mass is within 3 Da of expected. **H.** ESI-MS trace for H4K12acK16ac, Calculated mass is within 3 Da of expected. **I.** ESI-MS trace for H4K5acK8acK12acK16ac, Calculated mass is within 3 Da of expected. **J.** Analytical HPLC traces collected using a C4-analytical column with a gradient of 0-100% MeCN, 0.1% TFA in Milli-Q water, 0.1% TFA over 30 min. Representative traces for each ligation with unique ligation junctions are shown.

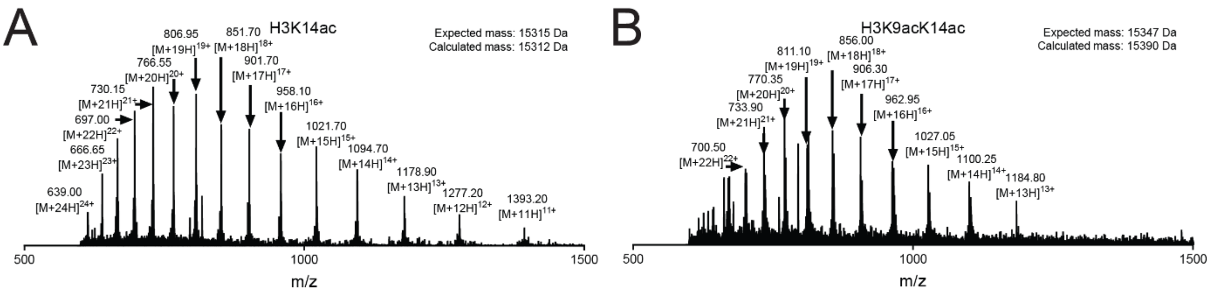


**Supplementary Figure 6: ESI-MS traces for modified histones expressed using an Amber stop codon suppression system. A.** ESI-MS trace for H3K14ac, Calculated mass is within 3 Da of expected. **B.** ESI-MS trace for H3K9acK14, Calculated mass is 33 Da larger than expected, however sequencing following MS/MS revealed the correct lysines were modified with N-acetyl groups. The observed mass difference was identified on fragments containing the two cysteines present in H3.1 and corresponded to an unidentified +16 Da modification.

**Supplementary Table 1: Proportions of histone modifications in native nucleosome library before and after pulldown with BRD4.** Proportions calculated from intensities for each modified peptide divided by the sum of all peptides for a family. Experiment was conducted with two biological replicates.

| | Input Proportions | | Pulldown Proportions | | Fold changes | |
|---|---|---|---|---|---|---|
| Modification | Input_1 | Input_2 | BRD4_1 | BRD4_2 | FC_1 | FC_2 |
| H3K4un | 0.78084 | 0.80541 | 0.79648 | 0.86656 | 1.02 | 1.08 |
| H3K9unK14un | 0.13386 | 0.26439 | 0.17572 | 0.21570 | 1.31 | 0.82 |
| H3K18unK23un | 0.86473 | 0.87976 | 0.71589 | 0.81539 | 0.83 | 0.93 |
| H3K27unK36un | 0.00519 | 0.02826 | 0.00292 | 0.03043 | 0.56 | 1.08 |
| H3K79un | 0.92940 | 0.95148 | 0.90707 | 0.93568 | 0.98 | 0.98 |
| H4K5unK8unK12unK16un | 0.78290 | 0.66907 | 0.65790 | 0.50216 | 0.84 | 0.75 |
| H4K20un | 0.59422 | 0.48450 | 0.43957 | 0.47761 | 0.74 | 0.99 |
| H3K9acK14un | 0.00094 | 0.00530 | 0.00263 | 0.00989 | 2.80 | 1.86 |
| H3K9unK14ac | 0.02233 | 0.01341 | 0.06462 | 0.01968 | 2.89 | 1.47 |
| H3K9me1K14ac | 0.01278 | 0.00508 | 0.02782 | 0.00601 | 2.18 | 1.18 |
| H3K9me2K14ac | 0.04910 | 0.00319 | 0.02254 | 0.01928 | 0.46 | 6.05 |
| H3K9me3K14ac | 0.03504 | 0.00126 | 0.00756 | 0.00797 | 0.22 | 6.31 |
| H3K18acK23un | 0.00511 | 0.00406 | 0.01091 | 0.00785 | 2.13 | 1.93 |
| H3K18unK23ac | 0.12638 | 0.11387 | 0.26484 | 0.17230 | 2.10 | 1.51 |
| H3K9acK14ac | 0.00086 | 0.00050 | 0.00396 | 0.00123 | 4.62 | 2.48 |
| H3K18acK23ac | 0.00154 | 0.00098 | 0.00779 | 0.00382 | 5.08 | 3.89 |
| H4K5ac | 0.00179 | 0.00473 | 0.00499 | 0.01104 | 2.79 | 2.33 |
| H4K8ac | 0.00497 | 0.00584 | 0.01052 | 0.01321 | 2.11 | 2.26 |
| H4K12ac | 0.00883 | 0.00479 | 0.01659 | 0.01097 | 1.88 | 2.29 |
| H4K16ac | 0.18289 | 0.28318 | 0.23541 | 0.33464 | 1.29 | 1.18 |
| H4K5acK8ac | 0.00021 | 0.00055 | 0.00103 | 0.00262 | 4.81 | 4.77 |
| H4K5acK12ac | 0.00120 | 0.00133 | 0.00441 | 0.00619 | 3.67 | 4.64 |
| H4K5acK16ac | 0.00101 | 0.00498 | 0.00415 | 0.01660 | 4.09 | 3.33 |
| H4K8acK12ac | 0.00077 | 0.00044 | 0.00270 | 0.00214 | 3.53 | 4.84 |
| H4K8acK16ac | 0.00363 | 0.00967 | 0.01227 | 0.02880 | 3.38 | 2.98 |
| H4K12acK16ac | 0.00769 | 0.00779 | 0.03117 | 0.03229 | 4.05 | 4.15 |
| H4K5acK8acK12ac | 0.00017 | 0.00021 | 0.00074 | 0.00109 | 4.39 | 5.13 |
| H4K5acK8acK16ac | 0.00023 | 0.00135 | 0.00146 | 0.00710 | 6.33 | 5.27 |
| H4K5acK12acK16ac | 0.00186 | 0.00337 | 0.00891 | 0.01816 | 4.79 | 5.39 |
| H4K8acK12acK16ac | 0.00125 | 0.00136 | 0.00573 | 0.00732 | 4.60 | 5.39 |
| H4K5acK8acK12acK16ac | 0.00060 | 0.00133 | 0.00202 | 0.00568 | 3.37 | 4.28 |
| H3K4me1 | 0.21916 | 0.19459 | 0.20352 | 0.13344 | 0.93 | 0.69 |
| H3K9me1K14un | 0.13386 | 0.26439 | 0.07791 | 0.08323 | 0.58 | 0.31 |
| H3K18me1K23un | 0.00224 | 0.00133 | 0.00056 | 0.00063 | 0.25 | 0.48 |
| H3K27me1K36un | 0.01479 | 0.03921 | 0.00577 | 0.03067 | 0.39 | 0.78 |
| H3K27me1K36me1 | 0.00906 | 0.01947 | 0.00353 | 0.01287 | 0.39 | 0.66 |
| H3K27unK36me1 | 0.00162 | 0.00765 | | 0.00668 | | 0.87 |
| H3K79me1 | 0.04090 | 0.01974 | 0.06035 | 0.03298 | 1.48 | 1.67 |
| H4K20me1 | 0.28333 | 0.48004 | 0.26380 | 0.34645 | 0.93 | 0.72 |
| H3K9me2K14un | 0.34287 | 0.34174 | 0.37406 | 0.34902 | 1.09 | 1.02 |
| H3K27me2K36un | 0.14414 | 0.17216 | 0.14270 | 0.19077 | 0.99 | 1.11 |
| H3K27unK36me2 | 0.00508 | 0.01351 | | 0.01408 | | 1.04 |
| H3K27me2K36me2 | 0.28237 | 0.22149 | 0.33800 | 0.24869 | 1.20 | 1.12 |
| H3K27me2K36me1 | 0.11579 | 0.12216 | 0.11307 | 0.12357 | 0.98 | 1.01 |
| H3K27me1K36me2 | 0.03387 | 0.03987 | 0.02399 | 0.04271 | 0.71 | 1.07 |
| H3K79me2 | 0.02970 | 0.02878 | 0.03258 | 0.03134 | 1.10 | 1.09 |
| H4K20me2 | 0.12245 | 0.03546 | 0.29664 | 0.17594 | 2.42 | 4.96 |
| H3K9me3K14un | 0.29690 | 0.22940 | 0.24317 | 0.28801 | 0.82 | 1.26 |

| | | | | | | |
|---|---|---|---|---|---|---|
| H3K27me3K36un | 0.17593 | 0.18419 | 0.17111 | 0.14192 | 0.97 | 0.77 |
| H3K27me3K36me1 | 0.09035 | 0.07584 | 0.06491 | 0.05470 | 0.72 | 0.72 |
| H3K27me3K36me2 | 0.10393 | 0.06059 | 0.13400 | 0.09645 | 1.29 | 1.59 |
| H3.3 K27unK36un | 0.01167 | 0.02158 | | 0.03442 | 0.00 | 1.59 |
| H3.3 K27me1K36un | 0.01838 | 0.02278 | | 0.02949 | | 1.29 |
| H3.3 K27me2K36un | 0.13298 | 0.17331 | 0.12391 | 0.17692 | 0.93 | 1.02 |
| H3.3 K27me2K36me1 | 0.17061 | 0.15378 | 0.14487 | 0.17561 | 0.85 | 1.14 |
| H3.3 K27me2K36me2 | 0.43511 | 0.42707 | 0.55962 | 0.40284 | 1.29 | 0.94 |
| H3.3 K27me3K36un | 0.11875 | 0.10871 | 0.09993 | 0.10564 | 0.84 | 0.97 |
| H3.3 K27me3K36me1 | 0.09894 | 0.07638 | 0.07168 | 0.05510 | 0.72 | 0.72 |

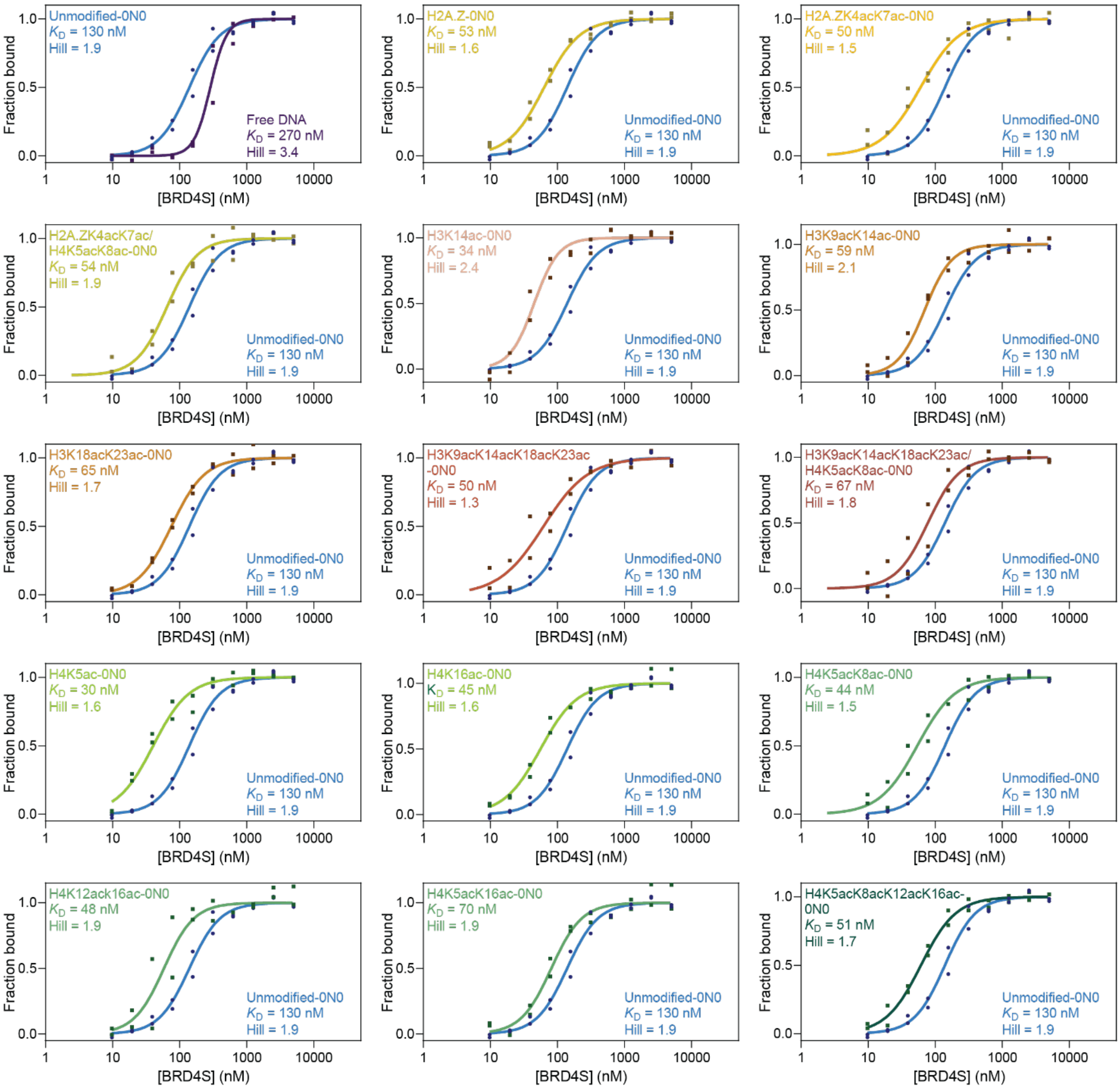


**Supplementary Figure 7: BRD4S binding to modified nucleosome library:** Technical duplicate MST assays for BRD4S binding to either unmodified or acetylated 0N0 Cy5-labelled nucleosomes. The SEM of all MST measurements is estimated to be 15% (see methods).

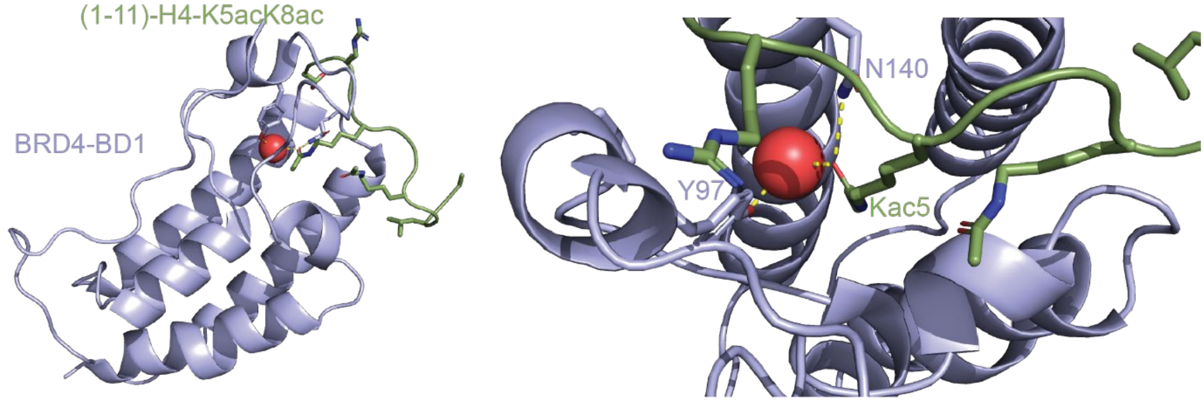


**Supplementary Figure 8: Structure of BRD4-BD1 in complex with H4K5acK8ac peptide.** PDB accession number 3UVW. Residues making direct or indirect polar contacts with H4K5ac are drawn with stick representation.

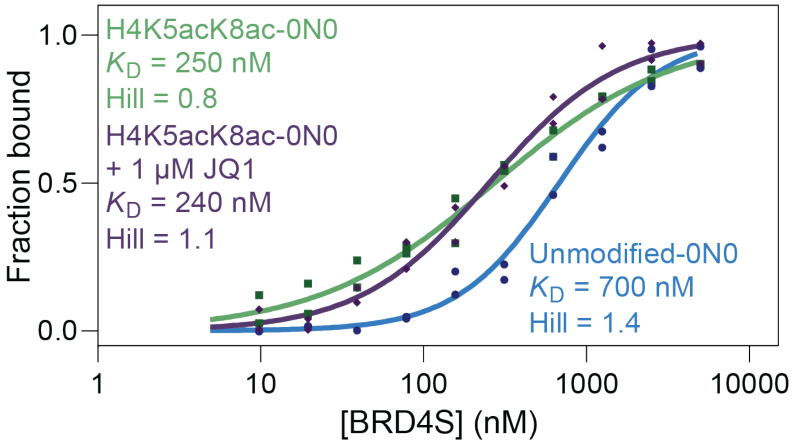


**Supplementary Figure 9: JQ1 inhibition at 150 mM NaCl.** MST titrations of BRD4S into unmodified and H4 acetylated nucleosomes, in the absence and presence of 1 µM JQ1 at 150 mM NaCl (technical duplicates). The SEM of all MST measurements is estimated to be 15% (see methods).
